# Genomic Loci Influencing Cue-Reactivity in Heterogeneous Stock Rats

**DOI:** 10.1101/2024.03.13.584852

**Authors:** Christopher P. King, Apurva S. Chitre, Joel D. Leal-Gutiérrez, Jordan A. Tripi, Alesa R. Hughson, Aidan P. Horvath, Alexander C. Lamparelli, Anthony George, Connor Martin, Celine L. St. Pierre, Thiago Sanches, Hannah V. Bimschleger, Jianjun Gao, Riyan Cheng, Khai-Minh Nguyen, Katie L. Holl, Oksana Polesskaya, Keita Ishiwari, Hao Chen, Leah C. Solberg Woods, Abraham A. Palmer, Terry E. Robinson, Shelly B. Flagel, Paul J. Meyer

**Affiliations:** Department of Psychology, University at Buffalo, Buffalo, USA; Clinical and Research Institute on Addictions, Buffalo, USA; Department of Psychiatry, University of California San Diego, La Jolla, USA; Department of Psychology, University of Michigan, Ann Arbor, USA; Department of Genetics, Washington University Saint Louis, St. Louis, USA; Department of Physiology, Medical College of Wisconsin, Milwaukee, USA; Department of Pharmacology and Toxicology, University at Buffalo, Buffalo USA; Department of Pharmacology, Addiction Science and Toxicology, University of Tennessee Health Science Center, Memphis, USA; Department of Internal Medicine, Molecular Medicine, Center on Diabetes, Obesity and Metabolism, Wake Forest School of Medicine, Winston-Salem, USA; Institute for Genomic Medicine, University of California San Diego, La Jolla, USA; Department of Psychiatry, University of Michigan, Ann Arbor, USA; Michigan Neuroscience Institute, University of Michigan, Ann Arbor, USA

## Abstract

Addiction vulnerability is associated with the tendency to attribute incentive salience to reward predictive cues; both addiction and the attribution of incentive salience are influenced by environmental and genetic factors. To characterize the genetic contributions to incentive salience attribution, we performed a genome-wide association study (GWAS) in a cohort of 1,645 genetically diverse heterogeneous stock (HS) rats. We tested HS rats in a Pavlovian conditioned approach task, in which we characterized the individual responses to food-associated stimuli (“cues”). Rats exhibited either cue-directed “sign-tracking” behavior or food-cup directed “goal-tracking” behavior. We then used the conditioned reinforcement procedure to determine whether rats would perform a novel operant response for unrewarded presentations of the cue. We found that these measures were moderately heritable (SNP heritability, *h^2^* = .189-.215). GWAS identified 14 quantitative trait loci (QTLs) for 11 of the 12 traits we examined. Interval sizes of these QTLs varied widely. 7 traits shared a QTL on chromosome 1 that contained a few genes (*e.g. Tenm4*, *Mir708*) that have been associated with substance use disorders and other mental health traits in humans. Other candidate genes (*e.g. Wnt11, Pak1*) in this region had coding variants and expression-QTLs in mesocorticolimbic regions of the brain. We also conducted a Phenome-Wide Association Study (PheWAS) on other behavioral measures in HS rats and found that regions containing QTLs on chromosome 1 were also associated with nicotine self-administration in a separate cohort of HS rats. These results provide a starting point for the molecular genetic dissection of incentive salience and provide further support for a relationship between attribution of incentive salience and drug abuse-related traits.

## Introduction

Addiction vulnerability is influenced by genetic and environmental factors. These factors are thought to include differences in cognitive and motivated behaviors, such as differences in the tendency to attribute incentive value to reward cues (Meyer, Ma, & Robinson, 2012; Saunders & Robinson, 2010; Versaggi et al., 2016), novelty-seeking (Beckmann et al., 2011; Belin et al., 2011; Molander et al., 2011), locomotor response to novelty (Gancarz et al., 2012, 2013; Piazza et al., 1989), and impulsivity (Belin et al., 2008; de Wit, 2009; Economidou et al., 2009; Perry et al., 2005). Thus a major avenue for understanding the genetics of addiction vulnerability is to delineate the genetic basis of these addiction-related traits.

Sensitivity to reward-paired stimuli is a particularly important addiction-related trait (Robinson & Flagel, 2009; Robinson et al., 2014), because incentive cues can instigate the craving and drug motivation that leads to relapse (Bailey et al., 2010; Carter & Tiffany, 1999; Shaham et al., 2003; Volkow et al., 2008). In rats, the incentive value of cues can be measured using a Pavlovian Conditioned Approach (PavCA) procedure, which measures individuals’ conditioned responses to a food-predictive reward cue. Some rats (“sign-trackers”; ST) show a strong tendency to approach and interact with cues that have become associated with a food reward, whereas others (“goal-trackers”; GT) show goal-directed responses in that they approach and interact with the food-delivery location (Boakes, 1977; Flagel et al., 2007; Hearst & Jenkins, 1974; Meyer, Lovic, et al., 2012). Thus, in sign-trackers, cues acquire incentive salience, as indicated by the extent to which cues elicit approach and become reinforcing (Robinson & Flagel, 2009). For incentive drug-cues, sign-trackers also show heightened responses to cocaine cues (Meyer, Ma, & Robinson, 2012), are sensitive to the ability of cocaine cues to support drug-taking (Saunders & Robinson, 2011), and motivate cocaine and nicotine-seeking (Pitchers et al., 2018; Saunders & Robinson, 2010; Versaggi et al., 2016). Sign-tracking is therefore an easily observed measure of cue-responsivity that predicts the effects of cues on several addiction related traits (Robinson et al., 2018).

Although there is substantial variability in the tendency to sign- or goal-track within outbred rat populations (Gileta et al., 2022; Hughson et al., 2019; King et al., 2021), few studies have examined the genetic basis for variation in the tendency to attribute incentive salience to cues (Dickson et al., 2015; Gileta et al., 2022). To address this knowledge gap, we conducted a genome-wide association study (GWAS) on PavCA to determine genetic underpinnings of sign- and goal-tracking in a large population (n=1,645) of heterogeneous stock rats (HS) rats (Hansen & Spuhler, 1984; Spuhler & Deitrich, 1984). HS rats were selected because of their high genotypic and phenotypic variability (Parker et al., 2014; Solberg Woods & Palmer, 2019), as well as the many complementary resources available for this population. We have used this HS population previously to compare sign- and goal-tracking to other drug-associated traits, including response to cocaine and cocaine cues (King et al., 2021).

## Materials and Methods

### Subjects

Subjects were NMcwi:HS rats (RRID:RGD_2314009; formerly known as N:NIH; N:NIH-HS; hereafter referred to as HS) that were shipped to the University at Buffalo from the laboratory of Dr. Leah Solberg Woods at Medical College of Wisconsin starting in 2015 until 2017. HS rats were originally established at the NIH by interbreeding eight inbred strains (ACI/N, BN/SsN, BUF/N, F344/N, M520/N, MR/N, WKY/N and WN/N; Hansen & Spuhler, 1984; Solberg Woods & Palmer, 2019). To maintain genetic diversity, HS rats were maintained using 48-64 breeding pairs per generation in conjunction with a breeding scheme that used kinship coefficients to minimize inbreeding. Wherever possible, no more than one male and one female per rat litter were shipped. This limited close relatives, which increases power in GWAS studies. This study used rats from generations 71 - 88.

Rats were shipped at approximately 33 days of age to the University at Buffalo, where they underwent 14 days of quarantine before being sent to the Clinical Research Institute on Addictions (CRIA) at the University at Buffalo. Rats of the same sex were pair-housed in plastic cages (42.5×22.5×19.25 cm) containing sawdust bedding (Aspen Shavings) in a temperature-controlled vivarium (22±1°C) with continuous access to water and food (Harlan Teklad Laboratory Diet #8604, Harlan Inc., Indianapolis, IN, USA). No environmental enrichment was provided. Rats underwent behavioral testing at the CRIA before being transferred to the University at Buffalo’s North Campus by laboratory animal facility staff (25-minutes by car). Traits measured during testing at the CRIA are being prepared for a separate publication and include tests for locomotor activity, light reinforcement, choice reaction time task, patch-depletion foraging test, and social reinforcement. Rats were acclimatized for a minimum of 7 days following transfer to North Campus, during which time they were handled daily. Rats were maintained on a reverse light/dark cycle at both CRIA and the University at Buffalo (lights off at 7:30am) and were tested a minimum of 1-hour following the onset of the dark cycle. PavCA testing began on average at PND162, range 140-204) in 16 batches, with each batch containing 7 groups of between 6-16 subjects. All studies were conducted according to the National Research Council (2003) “Guidelines for the Care and Use of Mammals in Neuroscience and Behavioral Research” and were approved by the University at Buffalo Institutional Animal Care and Use Committees.

### Procedure and Apparatus

#### Pavlovian Conditioned Approach (PavCA)

Rats were tested in 16 modular testing chambers (20.5×24.1 cm floor area, 29.2 cm high; MED-Associates Inc., St. Albans, VT) that were housed in custom built enclosures to attenuate external light and sound and were outfitted with fans that provided ventilation and background noise (A&B Display Systems, Bay City, MI). During PavCA, rats learned the association between presentation of a ∼45 mg banana-flavored food pellet and a conditioned stimulus (CS) (a backlit lever-CS) over 5 sessions. First, prior to testing, rats were exposed to the flavored food pellets in their home cage for two days (∼25 pellets per day; Bio-Serv, Flemington, NJ, #F0059). Next, rats underwent a single day of food cup training, in which subjects underwent a 5-minute chamber habituation before receiving 25 pellets delivered into an infrared photobeam equipped food-cup, or “magazine”, on a variable interval (VI)-30s (1-60s range) schedule. Details about the testing apparatus and equipment can be found on Open Behavior (https://edspace.american.edu/openbehavior/project/pavca/).

Rats then received five daily conditioning sessions in which they received 25 lever-food pairings, with each food pellet delivery preceded by the presentation of a retractable backlit lever for 8s. During testing, chambers were illuminated by a red light (27 cm high) on the back wall of the testing apparatus, and retractable levers were situated on either the left or right side of the food cup (2cm length, 6cm above floor). Lever presses and head entries into the food cup had no programmed consequences. Each lever-food pellet trial was separated on a VI-90 schedule (30-150s range) such that sessions lasted an average of 37.5 minutes. A summary of the testing procedure is available on protocols.io (https://dx.doi.org/10.17504/protocols.io.x54v9yjx4g3e/v1; Hannan, 2022).

#### Conditioned Reinforcement (CRf)

CRf was conducted the day after the final Pavlovian conditioning session and was used to assess the effectiveness of the lever-CS to reinforce a new instrumental response. Testing was conducted in the same chamber as PavCA, although the devices inside the chamber were organized differently. Specifically, the retractable lever was moved to the center of the instrument panel, and the food-cup was removed entirely. On either side of the lever were two nose poke ports with head-entry detectors. Each of the two nose poke ports were assigned as either active or inactive. Entries into the active port resulted in a 3-s delivery of the lever-CS. Responses in the inactive port had no programmed consequences. Sessions lasted 40 minutes. All data for PavCA and CRf were collected using the Med-PC IV software package (version 4.2, build 56).

### Behavioral Measures

We examined lever- and food-cup directed behavior during Pavlovian conditioned approach (PavCA). Approach to the lever was operationalized by incidental lever deflections (i.e. lever contacts) whereas approach to the food-cup was operationalized as food cup entries (food-cup contacts) during each of the 25 trials. Food cup entries during the inter-trial-interval period were also recorded. For each trial, the latency to deflect the lever or enter the food cup were also recorded. Previously, we have used these measures to calculate a general tendency to engage with the lever (“sign-tracking”) or food cup (“goal-tracking”) by calculating the PavCA index (Meyer, Lovic, et al., 2012). The index contains several calculated measures: 1) The *probability differential* of contact with the lever versus food-cup during each CS period (average probability of a lever contact on a given CS trial – average probability of a food-cup contact on a given CS trial), 2) the *response bias* directed towards either the lever or the food cup (# lever contacts - # food-cup contacts / # lever + # food-cup contacts), and finally 3) a *latency score* across trials to initiate contact with either the lever or food-cup (food-cup latency – lever latency / 8). The *PavCA index* was computed by averaging these three measures, yielding a value from −1 to 1, with −1 reflecting exclusive tendency to goal-tracking and 1 reflecting exclusive tendency to sign-track.

For Conditioned Reinforcement (CRf) the primary measures were total active and inactive responses, total earned lever reinforcers, total lever deflections, and an incentive value index ((responses in active port − responses in inactive port) / lever contacts). We chose to separately examine total lever deflections and lever deflections corrected for total responses, because we have previously shown that these measures are more strongly correlated to the PavCA index (Hughson et al., 2019; King et al., 2021).

There was substantial variability in tendency to sign- and goal-track during PavCA in both males and females, and the tendency to sign-track was strongly associated with the subsequent reinforcing value of the lever during CRf (r^2^ = 0.39 and 0.48 for males and females, respectively) as described previously (King et al., 2021). The behavioral analyses of these two tasks are described in detail in King et al., (2021) and so for brevity, we do not present these data here. Selected measures from the two tasks, described below, were used to examine genetic loci associated with tendency to attribute incentive salience to reward cues (i.e. sign-track).

### Selection of measures

We focused on a battery of 12 measures that reflected key terminal (i.e., session 5) indicators of sign- and goal-tracking. Specifically, all session 5 PavCA measures included **1)** overall terminal PavCA index, **2)** lever CS latency, **3)** lever CS contacts, **4)** lever CS probability, **5)** food-cup CS contacts, **6)** food-cup CS latency, **7)** food-cup CS probability, **8)** response bias, and **9)** intertrial interval (ITI) contacts with the food-cup. For conditioned reinforcement, we focused on **10)** active–inactive ratio, **11)** subsequent number of lever presses, and **12)** overall incentive value index. We have shown previously that this set of 12 behaviors are stable by the end of conditioning, and most directly related to the sign- and goal-tracking phenotypes (King et al., 2021).

### Tissue collection and Genotyping

Upon completion of behavioral testing, spleens were collected from each rat and then transferred to the University of California San Diego for genotyping using genotyping-by-sequencing (Gileta et al., 2020; Parker et al., 2016), which produced 3,400,759 single-nucleotide polymorphisms (SNPs) with an estimated error rate <1%. All coordinates are based on the Rnor_6.0 assembly (Accession number GCA_000001895.4) of the rat genome. The sex chromosomes (X and Y) and mitochondria were not genotyped.

### Phenotypic and genetic correlations, heritability estimates

To address the non-normal distribution of several of the traits (phenotypic distributions are available in *Supplementary Materials)*, and to remove potential sex differences, each trait was quantile-normalized separately for males and females. The quantile normalization procedure randomly breaks ‘ties’ such that when two or more individuals have identical values, they are assigned different values in the output. Other covariates, including age and batch number were examined for each trait, and regression was used to correct for covariate effects if they explained more than 2% of the variance. The resulting residuals were quantile-normalized again and then used for GWAS. The Spearman test was used for phenotypic correlations. SNP heritability estimates were obtained using the REML method, and genetic correlations between traits were computed through bivariate GREML analysis, both performed with GCTA (Lee et al., 2012; Yang et al., 2011).

### Genome-Wide Association Analysis

To perform GWAS we used a linear mixed model, as implemented in GCTA (Lee et al., 2012; Yang et al., 2011), using all SNP genotypes to create a genetic relatedness matrices (GRM) which accounted for the complex familial relationships that are characteristic of laboratory populations like the HS rats. We used the Leave One Chromosome Out (LOCO) method to avoid proximal contamination (Cheng et al., 2013; Gonzales et al., 2018). Using permutation for a genome-wide alpha of <0.05, the significance threshold was -log(p)> 5.95, and for 10%, it was -log(p) > 5.67. Because all traits were quantile normalized, the same threshold can be used for all traits (Cheng & Palmer, 2013). QTLs were identified by scanning each chromosome to determine if there was at least one SNP that exceeded the permutation-derived threshold of−log10(p) > 5.95 which corresponds to a genome-wide threshold of p<0.05. To avoid spurious results, we required that each QTL be supported by at least one additional SNP within 0.5 Mb that had a p-value within 2 − log10(p) units. To detect multiple significant loci on the same chromosome, we initially selected the most significant SNP on a given chromosome and then used that SNP as a covariate and performed a second scan of the same chromosome to determine whether there was a second significant and conditionally independent QTL on the same chromosome. If necessary, we would have continued to repeat this process until no further significant QTLs were detected on the chromosome in question. This procedure was performed for each autosome. Because it is difficult to plot the results of these conditional analyses, Manhattan plots show the product of the initial scan prior to any conditional analysis.

## Results

Multiple measures of sign- and goal-tracking were collected across the five days of conditioning. We chose to focus on the final day of conditioning (session 5), which most directly reflects the stable sign- and goal-tracking phenotype. For CRf we focused on three key measures of the reinforcing value of the lever. Detailed results for all other measures across the 5 conditioning sessions are reported in *Supplementary Material*. Note that the term “magazine” is used to refer to the food-cup in the supplementary material.

### Genetic correlations

The genetic correlation analysis of PavCA and CRf measures indicated significant shared genetic influence on a pair of behaviors (Lee et at. 2012). To examine the genetic relatedness among PavCA and CRf, phenotypic and genetic correlations (rG) for the set of behavioral measures were computed (**Figure 1**). Notably, two measures reflecting the attribution of incentive salience to the reward cue, “PavCA: Lever Presses” and “CRf: Incentive Value Index”, show a very high correlation (rG = 0.954), suggesting a shared genetic basis. Some measures show inverse phenotypic relationships between similar measures of sign-tracking, such as “PavCA: Lever CS Latency”. As a result, Lever CS latency has a negative genetic correlation with “PavCA: Lever Presses” (rG = −0.849). The “PavCA: Terminal Index” and related measures also exhibit strong positive correlations, underscoring their genetic relatedness. However, general activity measures like “PavCA: Food-Cup ITI” show weaker correlations, highlighting distinct genetic influences on other behaviors. Overall, the results reveal a shared genetic architecture underlying behavioral measures reflecting sign-tracking.

**Figure 1:**
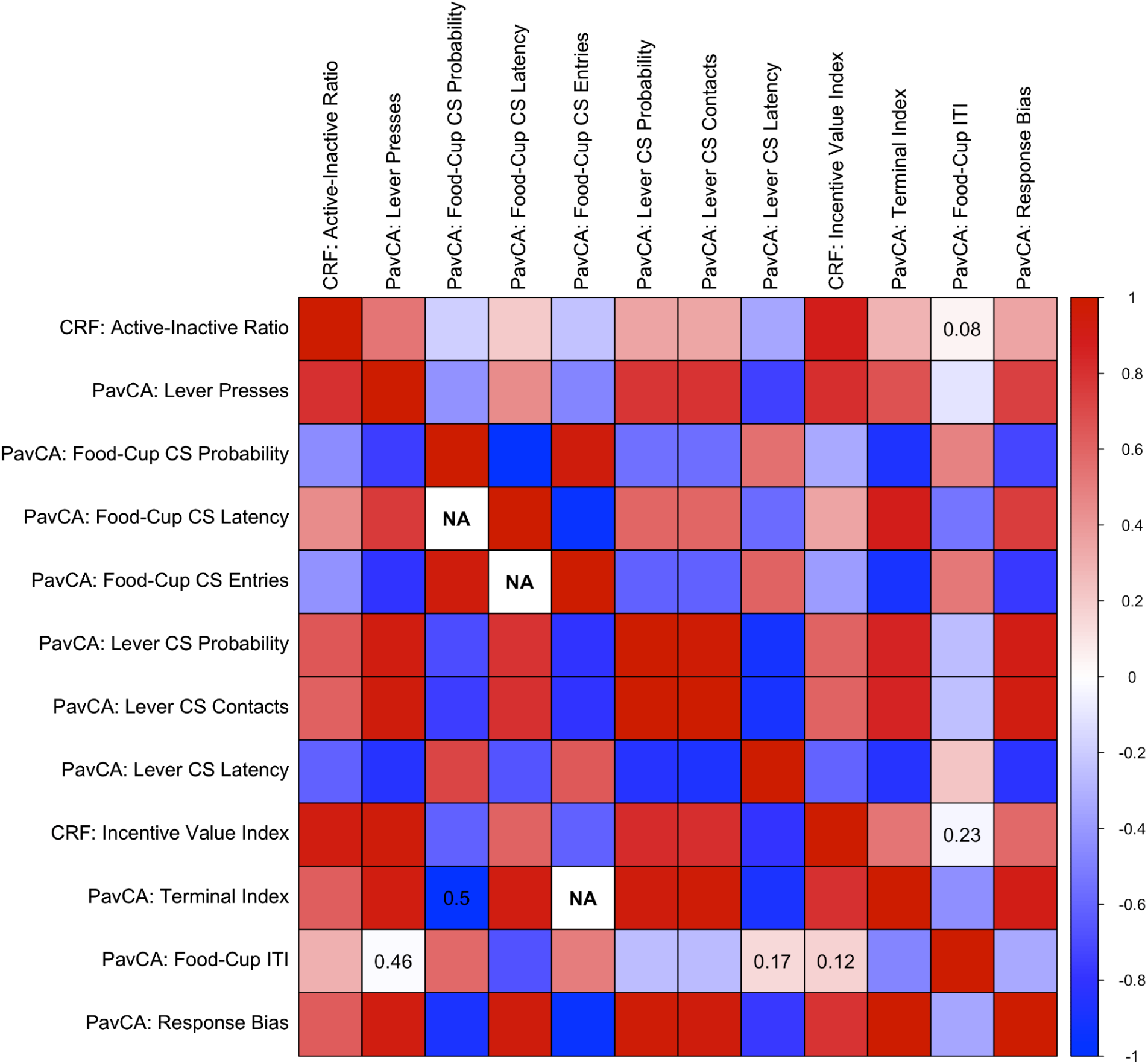
Phenotypic and genetic correlations for key sign- and goal-tracking measures. Phenotypic correlations between measures from day 5 are shown in the top-right triangle, and genotypic correlations are shown on the lower-left triangle. Red and blue squares reflect positive and negative correlations, respectively. The strongest genetic correlations were between sign- and goal-tracking measures, respectively, with weaker correlations occurring with inter-trial interval food-cup entries. All correlations were significant (*p* < 0.05) except where p-values are numerically indicated. ‘NA’ values denote genetic correlation pairs that were excluded due to non-invertible variance-covariance matrices, likely reflecting multicollinearity or insufficient variation.

### PavCA and CRf show modest heritability

Next, we examined SNP heritability. In general, heritability was moderate across the different measures of PavCA and CRf, with heritability estimates for the different sign- and goal-tracking traits during PavCA ranging from 0.215 ± 0.04 (latency to lever contact) to 0.107 ± 0.02 (probability of food-cup entry) (**Table 1a**). Heritability estimates are shown in descending order for the various measures, with the strongest heritabilities reflecting measures related to terminal sign-tracking. CRf heritability also showed similarly modest values, with the highest relating to traits most directly reflecting lever-directed sign-tracking behavior during CRf (**Table 1b**).

**Table 1:**
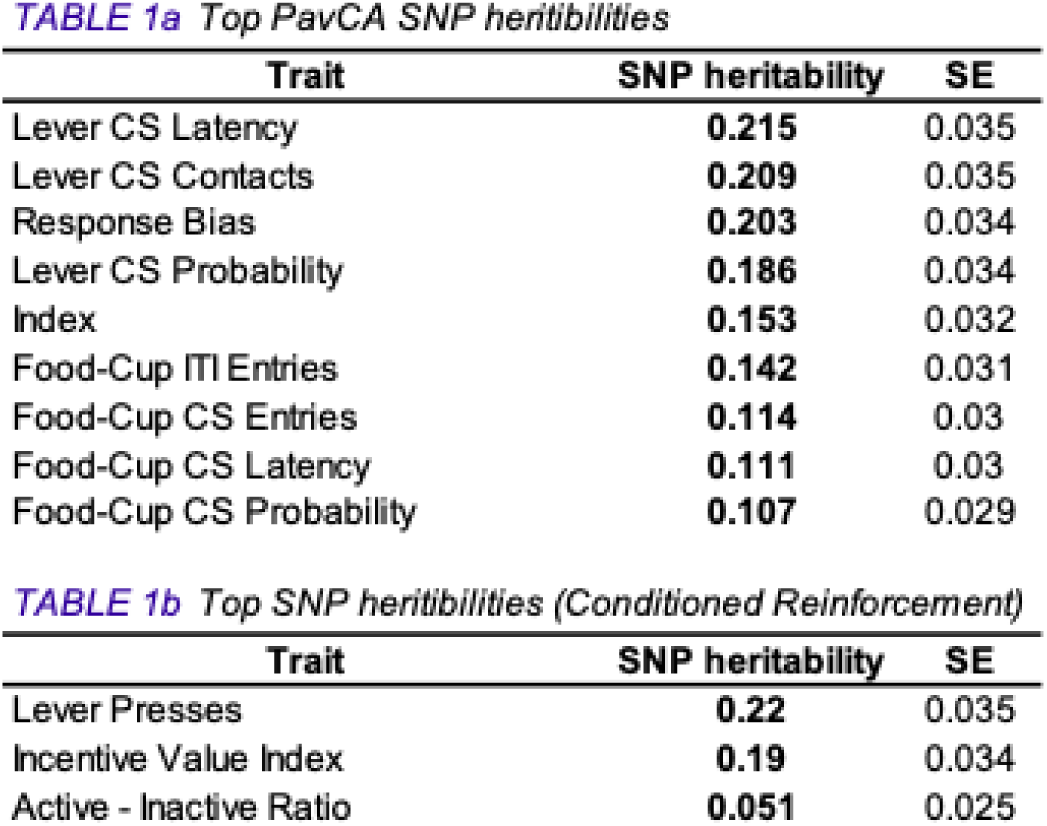
SNP heritability estimates for sign- and goal-tracking. **a)** The highest SNP heritability estimates tended to reflect measures of sign-tracking at the end of training relative to measures of goal-tracking. **b)** Similarly, conditioned reinforcement also showed modest SNP heritability, with the most heritable traits reflecting measures that directly assess lever-directed sign-tracking behavior (lever presses and overall incentive value index).

### Identification of multiple GWAS hits

We next performed a GWAS to identify specific genetic loci that were significantly associated with the tendency to sign- and goal-track. At least one QTL was identified for 11 of the 12 measures, with some measures being associated with more than one QTL and some QTLs being associated with more than one trait, such that a total of 6 different QTLs were identified for the 12 measures (**Table 2a**). Two of the three CRf QTLs overlapped with PavCA QTLs (**Table 2b**).

**Table 2:**
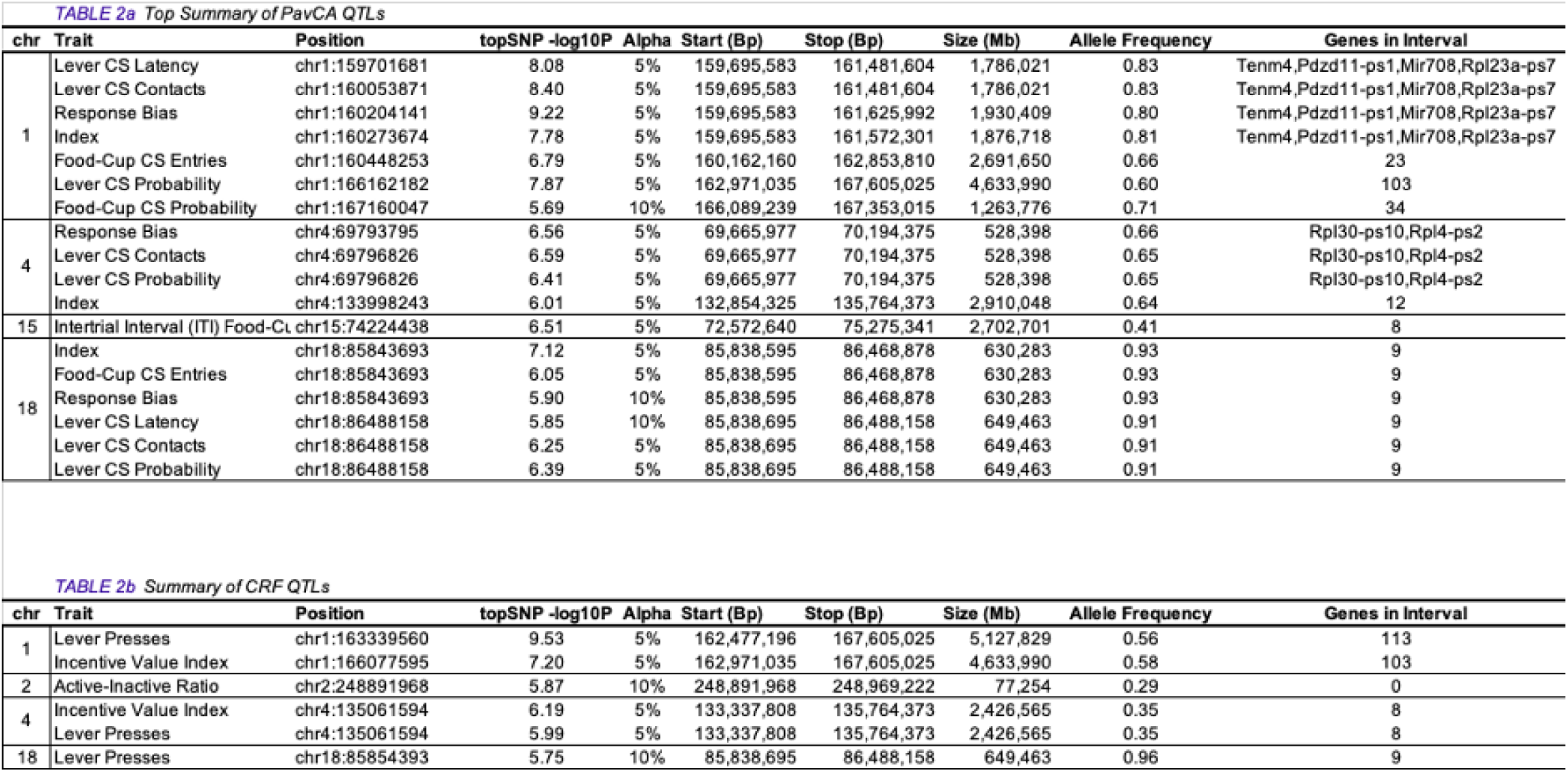
QTLs identified for PavCA and CRf. Shown are top SNPs and the associated QTL across the 12 traits examined in this GWAS. QTLs for a) PavCA and b) CRf are sorted by the chromosomal location of the top SNP position. Regions are blocked by chromosome. Top SNPs during PavCA largely reflected sign-tracking measures with QTLs identified on chromosomes 1, 4, and 18. Top SNPs during CRf also included overlapping regions of chromosomes 1 and 18 for lever-directed behavior. Identified QTLs varied in the number of genes contained in that interval, in some cases reflecting gene rich areas with as many as 113 genes. *Mir708*, microRNA 708; *Tenm4*, teneurin transmembrane protein 4; *Pdzd11-ps1*, PDZ domain containing 11, pseudogene 1; *Rpl23a-ps7*, ribosomal protein L23A, pseudogene 7. All coordinates are based on the Rnor_6.0 assembly of the rat genome.

Many loci had pleiotropic effects such that the same locus was implicated in two or more measures; the most notable example was on chromosome 1 (**Table 2a, 2b**). Three additional loci on chromosomes 4 and 18 were identified for both PavCA and CRf. The number of genes identified in the various QTLs ranged from 2 to 113 genes (full list of identified QTLs and Manhattan plots reported in *Supplementary Materials*).

**Figure 2:**
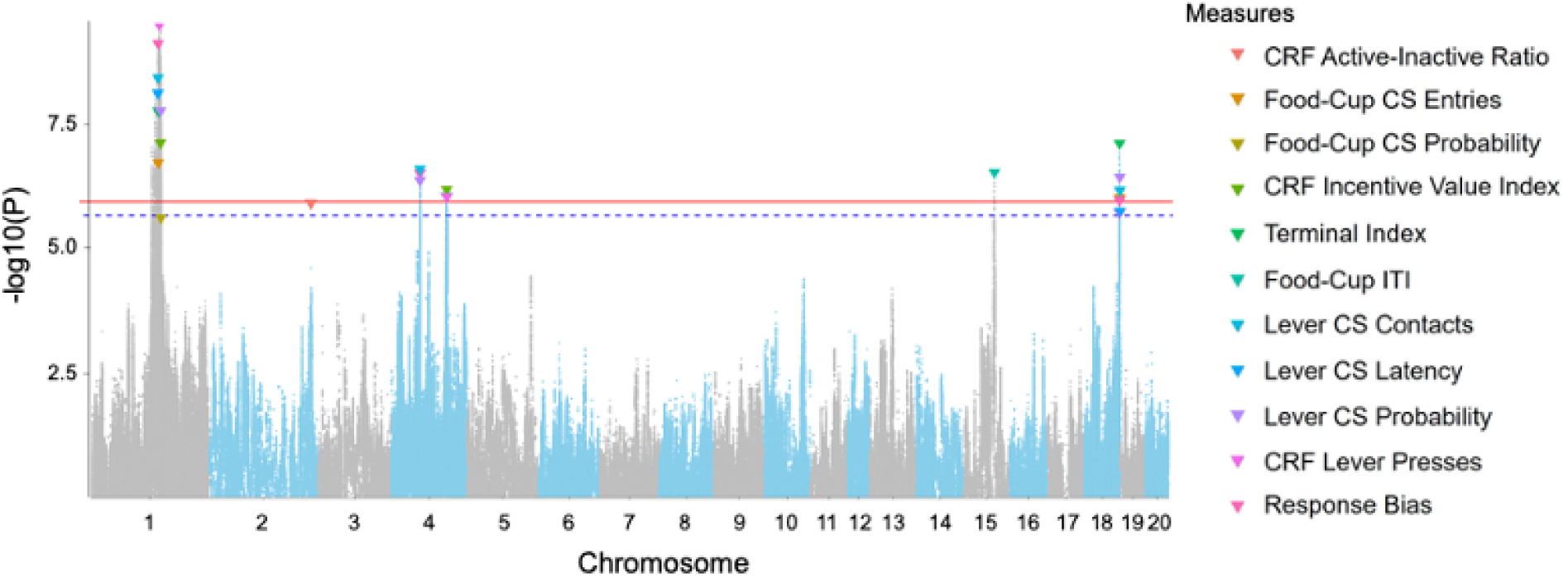
Porcupine plot for selected PavCA and CRf measures. Combined Manhattan plots from the genome-wide association study (GWAS) data for 11 traits with significant SNPs for sign- and goal-tracking. Chromosomal distribution of all P values (-log_10_ P values) is shown, and top SNPs are indicated by colored triangles. Cutoffs for genome-wide alpha of < 0.05 (-log_10_(p) > 5.95) is shown as the solid red line, and alpha < 0.10 (-log_10_(p) > 5.67) is shown as the dotted blue line. The largest cluster of SNPs was located on chromosome 1. For PavCA sign-tracking traits, more than one top SNP was identified in overlapping regions on chromosomes 1, 4, and 18. CRf lever-directed behavior also showed QTL overlap with similar regions of chromosome 1 and 18, relative to PavCA.

The chromosomal locations for identified regions of interest are shown below as a porcupine plot (**Figure 2**). Traits related to sign-tracking showed generally similar patterns, with overlap of the identified loci occurring on chromosomes 1, 4, and 18; These similar results partially reflect the high correlations among measures (**Figure 1**). Specifically, measures of sign-tracking during PavCA (response bias, lever latency, lever contacts) and CRf (incentive value index, lever presses) overlapped at each of these three regions.

### Candidate Gene Identification

The number of genes within each QTL varied from 2 to 113. We used several criteria to narrow down the list of candidate genes. For regions that contained multiple genes, we examined coding variants with moderate- to high-impact on protein function. We also examined genes that were differentially expressed in the central nervous system (CNS), or had functional relevance from the literature (*i.e.* also identified in human GWAS on psychiatric traits).

Figure 3 shows two representative regional association (“LocusZoom”) plots for QTLs on chromosomes 1 and 18. Additional LocusZoom plots are provided in the *Supplementary Materials.* One QTL that contained four genes was identified for 4 behavioral measures on chromosome 1 (Figure 3A). Two of the genes in this interval were *Tenm4* and *Mir708*. *Tenm4* expression is regulated by *Mir708* and has been previously identified as a candidate genetic component for psychiatric disorders (Lotan et al., 2014). There were also two other recently annotated pseudogenes (*Pdzd11*-*ps1*, *Rpl23a*-*ps7*) which are not well-characterized. A separate QTL on Chromosome 1 contained more than 100 genes (some of which are discussed later). The QTL identified on chromosome 18 for 7 different measures (Figure 3B) contained 9 genes (4 shown: *Socs6, Rttn, Cd226, Dok6*) (not shown in LocusZoom plot: *Pclaf-ps2*, PCNA clamp associated factor, pseudogene 2; *Chn3,* chimerin 3; *LOC689116* (*Ncbp2,* nuclear cap binding protein subunit 2); *LOC100362807*; *Snapc5-ps1*), snRNA-activating protein complex subunit 5, pseudogene 1). Two of these genes code for protein products which are involved in functionally regulating tyrosine kinase expression (SOCS6; Kabir et al., 2014) and binding to tropomyosin-related kinase receptors, influencing nervous system development (Dok6; Li et al., 2010). Table 2 provides a numerical summary of each region’s gene set size; detailed information can be found in *Supplementary Materials* with information such as strain distribution patterns (SDPs), LocusZoom plots, and the full list of genes for each interval.

**Figure 3:**
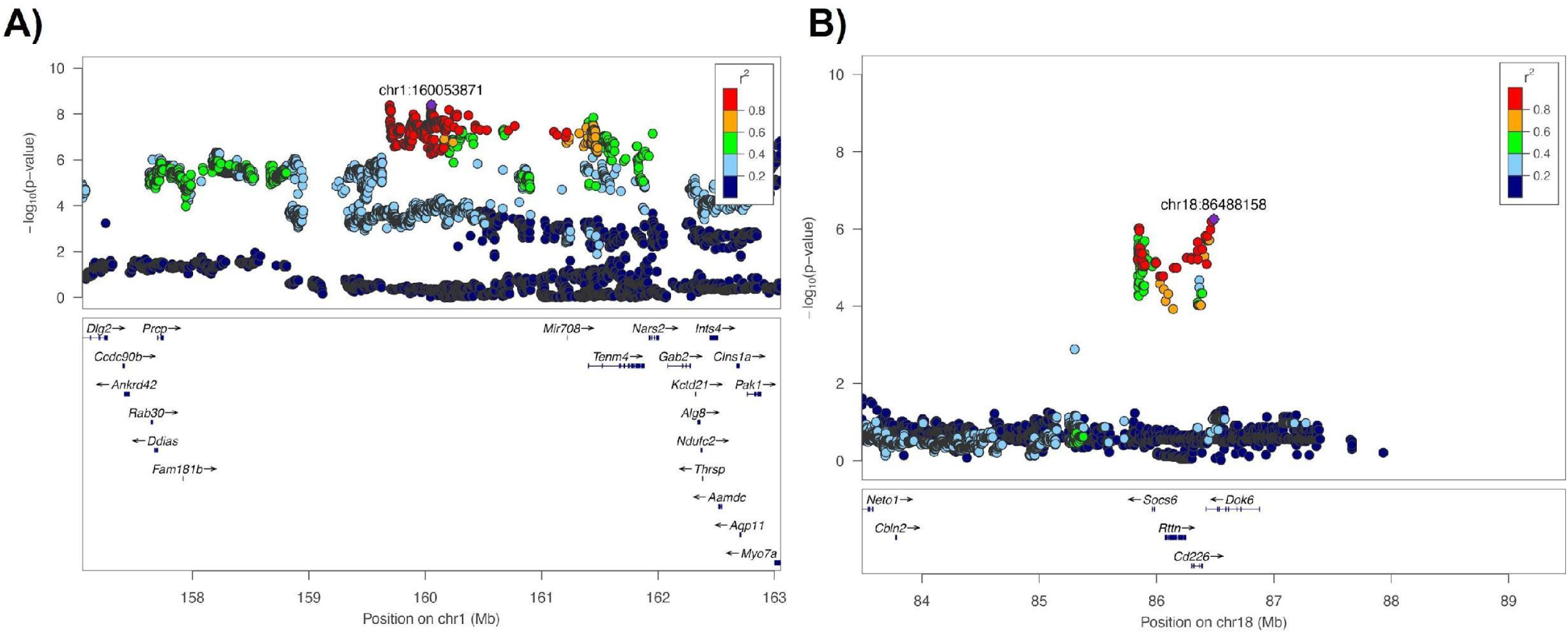
LocusZoom regional association plots of two top QTLs identified on chromosomes 1 and 18. The x-axis shows chromosomal position. The single-nucleotide polymorphism (SNP) with the lowest p-value is shown and labeled in purple (“top-SNP”). Color of dots indicates the degree of linkage disequilibrium (LD) of other SNPs relative to the top-SNP. The bottom half of each panel shows the genes in a particular region as annotated by the reference sequence. Panel **A)** shows genes contained in the QTL identified by the top SNP, two of which are named genes with known interactions (*Tenm4, Mir708).* Eight measures were associated with a QTL on chromosome 18 **B)**, which contains four genes (*Socs6*, suppressor of cytokine signaling 6; *Rttn*, rotatin; *Chn3*, chimerin 3; *Dok6*, docking protein 6).

To identify candidate genes in larger gene-rich regions, we looked for coding variants with potentially damaging effects on protein coding. For example, on chromosome 1, a 4.6MB region was identified with 103 genes. A total of 11 genes with moderate coding variants were identified across the entire set of QTLs (*Usp35, Alg8, Tsku, Serpinh1, Atg16l2, Art2b, LOC102549471, Chrna10, Nup98, Shq1, Numa1*). A subset of these genes are shown (**Table 3a**). We examined candidate genes using the PubMed and GWAS catalog mining tool GeneCup (Gunturkun et al., 2022), to probe for results from previous omics and gene-function studies.

**Table 3:**
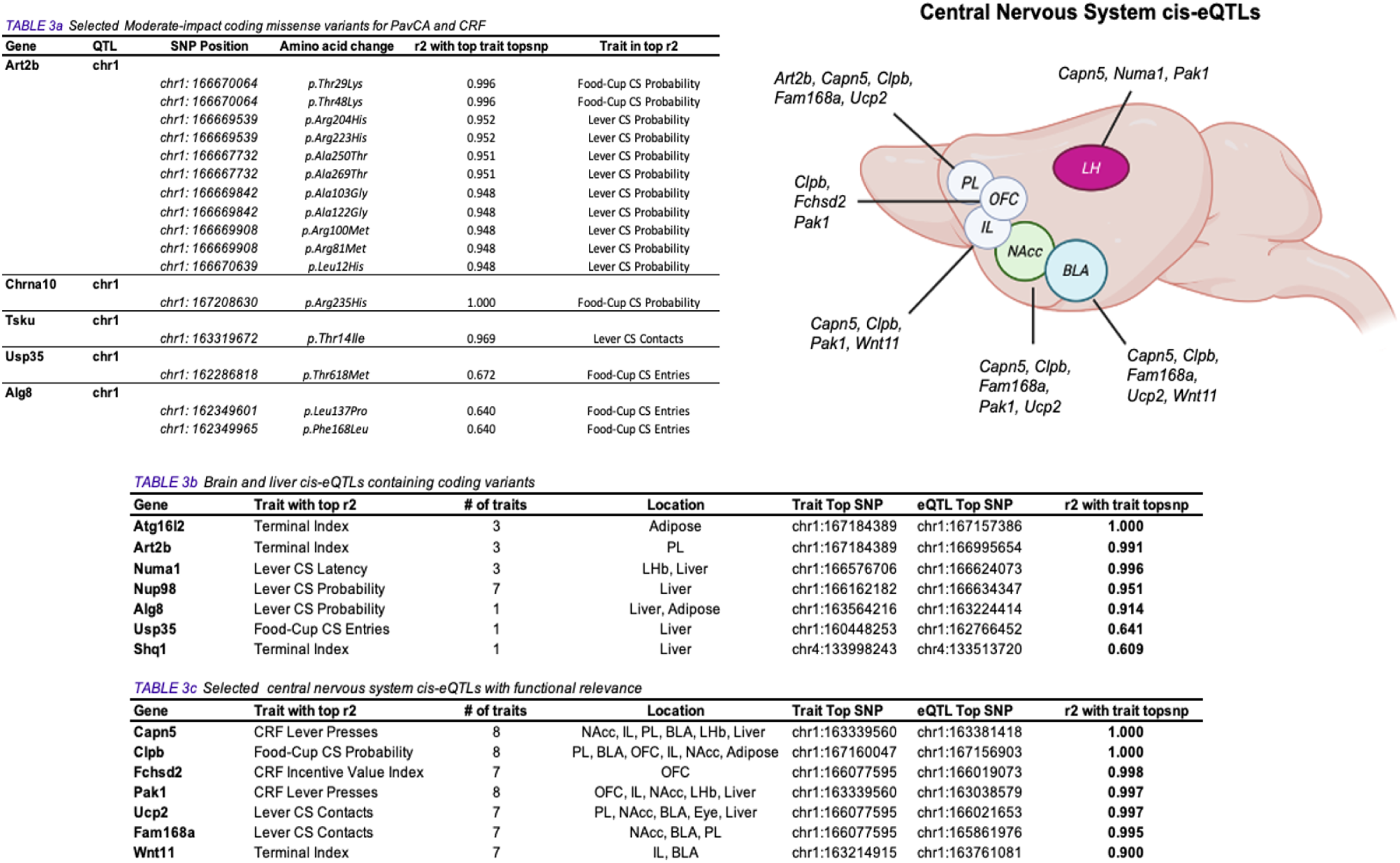
Coding variant impact chart and cis-eQTLs for candidate genes. **A)** Coding variants with moderate impact were mostly found to be located on chromosome 1, including those influencing sign-tracking traits. All genes resulted in missense coding substitutions. Traits associated with coding variants in highest association (r^2^) with the top-SNP or functional relevance are shown. **B)** Eight genes with coding variants also had heritable differences in expression (cis-eQTL) in various tissues; many based on expression in the liver. **C)** Other functionally-relevant genes highly correlated with measures of sign-tracking had cis-eQTL throughout the central nervous system. Summarized cis-eQTL in the brain are shown in the top right of the figure (created with BioRender.com). *Art2b,* ADP-ribosyltransferase 2b; *Chrna10,* cholinergic receptor, nicotinic, alpha polypeptide 10; *Tsku,* tsukushi, small leucine rich proteoglycan; *Usp35,* ubiquitin specific peptidase 35; *Alg8,* alpha-1,3-glucosyltransferase; *Atg16l2*, autophagy related 16-like 2; *Numa1*, nuclear mitotic apparatus protein 1; *Nup98*, nucleoporin 98 and 96 precursor; *Shq1,* SHQ1, H/ACA ribonucleoprotein assembly factor; *Capn5,* calpain 5; *Clpb,* caseinolytic mitochondrial matrix peptidase chaperone subunit B; *Clpb,* ClpB family mitochondrial disaggregase; *Fchsd2*; FCH and double SH3 domains 2; *Pak1*, p21 (RAC1) activated kinase 1; *Fam168a,* family with sequence similarity 168, member A; *Ucp2,* uncoupling protein 2; *Wnt11,* Wnt family member 11.

### Local genetic effects on gene expression

In addition to coding polymorphisms, we also considered expression quantitative trait loci or (eQTLs), which are loci that confer heritable differences in gene expression. The eQTL data we used are available at www.ratGTEx.org (Munro et al 2022). When a behavioral QTL and an eQTL are located nearby and are in strong linkage disequilibrium, it is possible that the behavioral differences (the behavioral QTL) is caused by the expression differences (the eQTL). In addition to liver, eye, and and adipose tissue gene expression, we identified central nervous system cis-eQTLs in the infralimbic cortex (IL), prelimbic cortex (PL), orbitofrontal cortex (OF), nucleus accumbens (NAcc), basolateral amygdala (BLA), and lateral habenula (LHb) that were colocalized with the QTLs for PavCA and CRf. Many cis-eQTL were in high association (r^2^ > 0.9) with more than one behavioral trait. These brain regions were selected due to their functional relevance across various addiction-related behavioral traits and because a recent study had identified numerous eQTLs in these intervals (Munro et al., 2022). There were 66 genes with cis-eQTL identified in this dataset, but for brevity only selected genes are shown based on whether they contained a coding variant (**Table 3b**) or were highly associated with the behavioral trait and functionally relevant (**Table 3c**).

Some genes with coding variants (**Table 3a**) may be of unique interest for complex behavioral traits, because they are expressed in the central nervous system, such as *Tsku* (Ito et al., 2010; discussed below). Each of the coding variant genes *Art2b, Tsku,* and the peripheral nicotinic receptor gene *Chrna10* (Lips et al., 2006) are also notable given their high association with the top-SNP in that QTL, which is consistent with them being the putatively causal SNPs. Notably, *Alg8,* which has been previously reported in another -omics study (Heinzman et al., 2019) contained multiple coding variant SNP substitutions as well as a cis-eQTL in the liver (**Table 3b**). The complete list of genes with both a coding variant and a cis-eQTL is shown in **Table 3b**. Among these, two are involved in cellular regulation mechanisms, including *Nup98* (Radu et al., 1995) and *Atg16l2* (Yang & Klionsky, 2010). Five coding variant genes reflected cis-eQTLs identified in the liver.

Many genes with cis-eQTLs did not contain coding variants, although for many of these eQTLs, the top eQTL SNP was in high linkage disequlibrium (LD as measured by r^2^) with the SNP most strongly implicated in the PavCA or CRf behaviors (**Table 3c**). For brevity, only those eQTL with an r^2^ above 0.9 are shown (Full set of eQTLs is available in the *Supplementary Materials*). However, a set of 7 genes (*Capn5, Clpb, Fchsd2, Pak1, Fam168a, Ucp2, Wnt11*) were identified in multiple brain areas across multiple instances of sign- and goal-tracking traits. This set of eight genes serves a variety of functions (discussed below). Additionally, *Tenm4*, one of few genes with better-characterized psychiatric relevance (Lotan et al., 2014), was identified separately as a differentially expressed eQTL in the prelimbic cortex RatGTEx, although there was no cis-eQTL identified in this GWAS.

### Phenome-Wide Association Study (PheWAS) Analysis

Whereas in GWAS, many individual SNPs are tested for their association with a single phenotypes, in a Phenome-Wide Association Study (PheWAS) a single SNP is tested for association with a collection of many traits, which is referred to as the “phenome” (Bush et al., 2016). This approach is useful, because PheWAS can identify genetic variants which are often known to exert multiple effects (Sanchez-Roige et al., 2023), and hence are pleiotropic.

We examined whether the genetic loci associated with the attribution of incentive salience were also associated with drug conditioning and other addiction-related behaviors collected in HS rats. Here, each 3MB window surrounding a top SNP for each identified QTL was tested for its association with a separate set of behavioral traits collected at each of three HS rat testing centers (University at Buffalo, University of Michigan, University of Tennessee Health Science Center) (**Table 4**). These traits included socially acquired nicotine self-administration (Wang et al., 2018), cocaine contextual conditioning (Hughson et al., 2019; King et al., 2021), sequential patch depletion (Gancarz et al., 2023; Ishiwari et al., 2023), light reinforcement (Gancarz et al., 2012, 2013; Ishiwari et al., 2023), locomotor response to novelty, reaction time, and social reinforcement (Ishiwari et al., 2023). **Table 4** shows key measures for each behavioral task, selected based on the relevance to the trait being measured (full list of PheWAS results for each task are available in *Supplementary Materials*)

**Table 4:** Phenome-Wide Association with drug conditioning and addiction related traits in HS rats. PheWAS tables show the association between addiction-related traits in other tests (“PheWAS Trait”) and measures from tests reported here in the GWAS for PavCA (“PavCA Trait in Top r^2^”). The chromosomal location for the association is shown as “Trait top-SNP”, and the strength of the association is shown as r^2^. In many cases, more than one PavCA QTL yielded a PheWAS association. The total number of PavCA measures in each QTL, and which PavCA categories those measures reflect are shown in the rightward columns. **A)** PheWAS are shown for two tasks measuring drug response to nicotine and cocaine. **B)** A separate cohort of rats that underwent an identical PavCA procedure at the University of Michigan were tested for PheWAS with the same measures for incentive salience described in this GWAS. **C)** PheWAS for multiple measures of behavioral regulation collected at the Clinical and Research Institute on Addictions (Buffalo, NY).

| PheWAS Trait | PavCA Trait in Top r2 | Trait topSNP | r2 | -log10P | dprime | # Measures | PavCA Behavioral Categories |
| --- | --- | --- | --- | --- | --- | --- | --- |
| **Nicotine Self-Administration** |  |  |  |  |  |  |  |
| Acquisition of Nicotine Self-Administration (Day 1 Responses) | Food-Cup CS Entries | chr1:160845434 | **0.844** | 4.295 | 0.993 | 6 | GT, ST, CRF, Overall |
| Reinstatement to Nicotine Seeking (Terminal Cue Responses) | CRF Incentive Value Index | chr1:163453166 | **0.289** | 4.141 | 0.918 | 2 | CRF, ST |
| Reinstatement to Nicotine Seeking (Terminal Cue Responses) | Food-Cup CS Entries | chr1:161226950 | **0.327** | 6.399 | 0.984 | 6 | GT, Overall, ST, CRF |
| Reinstatement to Nicotine Seeking (Terminal Cues Earned) | Food-Cup CS Entries | chr1:160554677 | **0.388** | 4.913 | 0.996 | 6 | GT, Overall, ST, CRF |
| Acquisition of Nicotine Self-Administration (Day 1-3 Responses) | Lever CS Contacts | chr1:160071477 | **1** | 4.052 | 1 | 5 | ST, Overall, GT |
| **Cocaine Contextual Conditioning** |  |  |  |  |  |  |  |
| Locomotion (Habituation) | CRF Active-Inactive | chr2:249632706 | **0.376** | 4.067 | 0.81 | 2 | CRF |
| Cocaine-induced locomotion (Velocity Day 3) | CRF Active-Inactive | chr2:250361653 | **0.013** | 4.197 | 0.965 | 2 | CRF |
| Locomotion (Habituation) | Response Bias | chr4:70595788 | **0.03** | 4.362 | 0.552 | 3 | CRF, ST, Overall |
| Cocaine-induced Sensitization (Change in Head-waving) | Food-Cup ITI Entries | chr15:72160751 | **0.06** | 4.211 | 0.522 | 1 | Non-specific |
| Cocaine-induced Stereotypy (Head-waving Day 7) | CRF Active-Inactive | chr2:251593694 | **0.192** | 4.079 | 0.624 | 2 | CRF |

| PheWAS Trait | PavCA Trait in Top r2 | Trait topSNP | r2 | -log10P | dprime | # Measures | PavCA Behavioral Categories |
| --- | --- | --- | --- | --- | --- | --- | --- |
| **Pavlovian Conditioned Approach (Michigan)** |  |  |  |  |  |  |  |
| Terminal Food-Cup CS Entries (GT) | Food-Cup CS Entries | chr18:85843691 | **0.943** | 4.229 | 1 | 7 | GT, Overall, CRF, ST |
| Terminal Food-Cup CS Probability (GT) | Food-Cup CS Entries | chr18:85843691 | **0.943** | 4.278 | 1 | 7 | GT, Overall, CRF, ST |
| Terminal Food-Cup CS Probability (GT) | CRF Active-Inactive | chr2:250881872 | **0.073** | 4.45 | 0.851 | 2 | CRF |
| **Conditioned Reinforcement (Michigan)** |  |  |  |  |  |  |  |
| Incentive Value Index (CRF) | Lever CS Contacts | chr18:86146075 | **1** | 6.666 | 1 | 7 | ST, GT, CRF |
| Lever Presses (CRF) | CRF Lever Presses | chr18:85859393 | **1** | 7.491 | 1 | 7 | ST, GT, CRF, Overall |

| PheWAS Trait | PavCA Trait in Top r2 | Trait topSNP | r2 | -log10P | dprime | # Measures | PavCA Behavioral Categories |
| --- | --- | --- | --- | --- | --- | --- | --- |
| **Patch-Depletion Foraging Test** |  |  |  |  |  |  |  |
| Water consumption rate (12s delay) | CRF Incentive Value Index | chr4:135452263 | **0.962** | 4.493 | 0.988 | 3 | CRF, Overall |
| Water consumption rate (6s delay) | Terminal Index | chr4:136848229 | **0.126** | 4.588 | 0.617 | 3 | CRF, Overall |
| Water consumption rate (No delay) | CRF Incentive Value Index | chr4:137509951 | **0.078** | 4.192 | 0.669 | 2 | CRF |
| Indifference Point (18s Delay) | Lever CS Contacts | chr4:67351289 | **0.001** | 4.043 | 0.062 | 3 | ST, Overall |
| **Light-Reinforcement** |  |  |  |  |  |  |  |
| Maximum Responses (Any 3 minute epoch) | CRF Active-Inactive | chr2:249349603 | **0.268** | 4.296 | 0.852 | 2 | CRF |
| Maximum Responses (First 3 minute epoch) | CRF Active-Inactive | chr2:249349603 | **0.268** | 6.316 | 0.852 | 2 | CRF |
| **Locomotor Response to Novelty** |  |  |  |  |  |  |  |
| Response to Novelty (Rearing) | Food-Cup CS Entries | chr18:86363830 | **0.752** | 5.954 | 0.995 | 7 | GT, ST, CRF, Overall |
| Response to Novelty (Locomotion) | Terminal Index | chr4:133631681 | **0.135** | 4.043 | 0.485 | 3 | CRF, Overall |
| **Reaction Time Task** |  |  |  |  |  |  |  |
| Change in Premature Responses Per Opportunity Across Time | Response Bias | chr4:69757343 | **0.757** | 4.115 | 0.986 | 3 | Overall, ST |
| **Social Reinforcement** |  |  |  |  |  |  |  |
| Time in Social Interaction Zone | Food-Cup ITI Entries | chr15:76781721 | **0.235** | 4.108 | 0.809 | 1 | Non-specific |
| Number of earned social reinforcers | Food-Cup ITI Entries | chr15:75366723 | **0.018** | 4.035 | 0.273 | 1 | Non-specific |

We found that loci associated with attribution of incentive salience overlapped with those associated with measures of drug response (**Table 4a**). As part of this process, we used a socially acquired adolescent nicotine self-administration protocol as described previously (Wang et al., 2018) where rats engaged in operant licking for infusions of nicotine. We examined two major features of nicotine-directed behavior: the acquisition of nicotine self-administration (day 1; day 1-3 responses) and reinstatement to nicotine seeking following extinction (terminal cue responses) as a measure of relapse. The strongest association (r^2^ > 0.84) was identified with initial responding for nicotine on chromosome 1. A weaker association was identified with reinstatement to nicotine seeking (r^2^: 0.29 – 0.39) similarly on chromosome 1.

In a separate task, cocaine contextual conditioning, rats receive repeated injections of cocaine in a designated cocaine “context” (Hughson et al., 2019). Here, sensitization to the locomotor activating and head-waving response is measured across 4 sessions. PheWAS for cocaine contextual conditioning revealed a region on chromosome 2 that was only modestly associated with head-waving on the final session of cocaine conditioning (r^2^ = 0.19) (**Table 4a**). Together, PheWAS for these two traits suggest a stronger association with nicotine response driven by loci on chromosome 1.

In addition to drug response, we used PheWAS to test the association between measures of sign- and goal-tracking with a separate cohort of HS rats that underwent an identical PavCA and CRf procedure at the University of Michigan (Hughson et al., 2019) (**Table 4b**). PheWAS yielded strong associations (r^2^: 0.73 – 0.93) for measures of goal-tracking during PavCA in the Michigan cohort, identifying overlapping regions on chromosomes 2 and 18. The association with measures of sign-tracking during CRf was particularly strong (r^2^ = 1) on chromosome 18, suggesting PheWAS can be used to validate chromosomal regions using separate but similarly phenotyped cohorts.

Finally, PheWAS was used to determine if genetic loci identified for PavCA overlapped with other measures of behavioral regulation (**Table 4c**) (Ishiwari et al., 2023). The QTL on chromosome 18 which was associated with food-cup CS entries also influenced the locomotor (rearing) response to novelty (r^2^ = 0.75). However, more complex behavior, including measures of foraging and impulse control, yielded PheWAS associations that varied widely by task and measure. For example, during a patch-depletion foraging test (Gancarz et al., 2023), water-deplete rats consume water in one of two “patches”, in which the amount of water available at a particular patch depletes over time. Switching patches is therefore adaptive in response to patch depletion that varies between subjects (Richards et al., 2013), but patch switching results in one of several experimenter-imposed delays. Rate of patch switching and consumption can therefore be used as a measure of foraging under different conditions. Incentive value index was associated with a QTL on chromosome 4 that was also associated with the rate at which rats maximized water consumption during this task, although the strength of the association was dependent on the length of imposed delay for switching patches (r^2^: 0.08-0.96). This PheWAS association is therefore dependent on task performance, specifically when the task is made most difficult by imposing a longer patch switching delay (12s).

## Discussion

This study is the first to use a large population of HS rats to identify genetic loci associated with the tendency to attribute incentive salience to reward cues, as measured by sign-tracking and conditioned reinforcement. Measures of sign-tracking were moderately heritable, and highly phenotypically and genetically correlated, suggesting common loci underlying individual variability in these measures. Among the ten unique loci identified using genome-wide association, there were multiple candidate genes, including previously identified SUD genes as well as genes not previously associated with SUD. Further, these identified QTLs reflected chromosomal regions that were significantly associated with other measures of other behavioral traits, including nicotine self-administration. These data demonstrate the utility of HS rats for the genetic mapping of complex behavioral traits. Some of the candidate genes identified are particularly promising targets with known functional or psychiatric relevance.

HS rats are useful for genetic mapping of small regions, in some cases the regions contain a small number of genes. However, other loci were relatively gene-rich regions, which made parsing the underlying candidate genes more difficult. Thus, we examined candidate genes in these QTLs by using several strategies: 1) Identifying genes with coding variants that are predicted to have moderate or large impacts on protein function, 2) identifying genes with corresponding eQTLs in relevant brain regions, and 3) identifying genes previously associated with psychiatric functions in other omics studies, including human GWASs. Thus, we highlight and discuss several of these genes in addition to presenting the report containing the full dataset of genes.

### Tenm4 and Mir708

One especially promising gene candidate identified inside a chromosome 1 QTL was *Tenm4*. The teneurins are surface-bound transmembrane glycoproteins conserved across species (Tucker et al., 2012; Tucker & Chiquet-Ehrismann, 2006), and are located in synapses with multiple functions including cell-adhesion (Araç & Li, 2019). *Tenm4,* which is expressed in the central nervous system (Zhou et al., 2003), is involved in functions such as axon guidance (Hor et al., 2015) and associated with disorders such as schizophrenia (Xue et al., 2019). Teneurin-4 interacts with a group of proteins involved in postsynaptic density function, which may be related to pleiotropic effects of *Tenm4* in multiple psychiatric disorders (Lotan et al., 2014).

Interestingly, the teneurins are well-situated to affect complex behavior through regulation of mood and cortiocotropin-releasing hormone (CRH)-mediated stress effects (Hogg et al., 2022; Woelfle et al., 2016; Woelfle et al., 2015) via cleavable teneurin C-terminal associated signaling peptides (TCAPs) (Lovejoy et al., 2006; Tucker et al., 2012; Tucker & Chiquet-Ehrismann, 2006) which work as extracellular soluble signaling proteins. The TCAPs (1-4) correspond to each of the teneurin 1-4 genes, and as such TCAP-4 is an interesting signaling peptide for future functional studies. In one earlier study, TCAP-1, which produces an anxiolytic effect (Al Chawaf et al., 2007), reduced stress-induced reinstatement to cocaine self-seeking (Erb et al., 2014). Although no behavioral studies have examined the role of TCAP-4, this may be one promising target for examining Tenm4 function with respect to behavior.

*Tenm4* is located in the same QTL as *Mir708.* In humans, *MIR708* is a microRNA contained in an intron of the protein coding gene, or mirtron, (Berezikov et al., 2007) for *TENM4.* Similar to *Tenm4*, *Mir708* is expressed in the brain and is differentially expressed across mesocorticolimbic circuitry in mice (Hamilton et al., 2014). Previous work suggests that dopamine and subcortical neural circuits are important in the attribution of incentive salience (Flagel & Robinson, 2017; Flagel et al., 2010; Haight et al., 2017), and therefore these two genes may work in concert to regulate forebrain function.

### Genes with coding polymorphisms

Several genes within QTLs contained putative coding variants. Several of these genes are expressed in the brain, which is consistent with them having a role in complex behavioral traits such as cue-responsivity. For example, *Tsku* is a member of the small leucine-rich proteoglycans (SLRPs) family (Deng et al., 2021) and has established functions in the CNS where it is crucial for commissure development (Ito et al., 2010) and has recently been characterized for roles in hippocampal neuronal development (Ahmad et al., 2020). Other genes such as *ALG8* have been identified by human GWAS of depression in smokers (Heinzman et al., 2019). *Chrna10* is another interesting candidate, which codes for nicotinic receptor alpha-10 subunit and is expressed in the ear (Fritzsch & Elliott, 2017; Taranda et al., 2009) and peripheral sympathetic nervous system (Lips et al., 2006). *Chrna10* has been previously associated with subjective response to nicotine (Ehringer et al., 2010) and dependence (Saccone et al., 2010). Other genes with coding polymorphisms and cis-eQTL are novel candidate targets, such as *Serpinh1, Atg16l2, Art2b,* and *Nup98,* which currently lack pre existing evidence implicating them in behavioral or psychiatric traits. Two of these genes are involved in cellular regulation, for example *Nup98* regulates transport of proteins into the cell nucleus (Radu et al., 1995) and *Atg16l2* regulates cellular autophagy (Yang & Klionsky, 2010). Among the coding variant containing genes, *Tsku, Chrna10* and *Alg8,* which have been previously implicated in central nervous system function, development, and psychiatric relevance may be attractive targets for their roles in complex behavior.

### Expression-QTL Genes

Using cis-eQTL analysis, we identified a group of eight genes that were all differentially expressed in regions of the brain, some of which have yet to be reported in the literature in relation to behavioral and nervous system function. To further investigate genes that had eQTLs that colocalized with behavioral QTL, we used GeneCup (Gunturkun et al., 2022) to retrieve psychiatrically and mechanistically relevant pre-existing literature.

*Capn5* is an attractive candidate gene given its particularly high association with multiple PavCA traits and five cis-eQTLs throughout the forebrain (NAcc, IL, PL, BLA, LHb). There are some experimental data indicating *Capn5* may be broadly involved in central nervous system function and expressed throughout the brain, including granule cells of the hippocampus, cerebellum, (Schaefer et al., 2017) and piriform cortex (Nakashima et al., 2019) where it functions enzymatically as a calcium-dependent protease (Bondada et al., 2021). The calpains have been implicated in neurodegenerative disorders (Araujo et al., 2008; Gao & Geng, 2013) but *Capn5* does not have well-characterized pathology-related mechanisms in the nervous system outside of the retina (Schaefer et al., 2016). The utility of these genes as genetic targets for regulation of behavior and cue-responsiveness will benefit from additional experimental and omics studies that examine functional and behavioral relevance to the various domains of psychological function.

*Pak1* is also notable for its strong association with four PavCA measures and four cis-eQTLs throughout the brain. Pak1 (p21 RAC1 activated kinase 1) is a kinase active in the central nervous system that is an effector of the family of Rac1 and Cdc42 GTPases, regulating neuronal morphology and synapses (Nikolic, 2008), axon migration and synaptic plasticity (Kreis & Barnier, 2009) and dendrite initiation (Hayashi et al., 2002). Further, PAK1 has been suggested to be involved in neurological disorders including schizophrenia (Jiang et al., 2017; Rubio et al., 2012) depression (Fuchsova et al., 2016), and other neurodegenerative diseases (Ma et al., 2012). It is therefore likely that *Pak1* is pleiotropic and broadly regulates complex behaviors.

*Ucp2* is another candidate influencing the response to food cues, given its expression in the NAcc and PL, high association with three measures of PavCA, and previously established roles in diet and food response. *Ucp2* (Uncoupling protein 2) codes for an uncoupling protein in the mitochondria that reduces ATP production with a role in energy balance (Fleury et al., 1997). *UCP2* is polymorphic in humans, which results in differential mRNA expression and association with obesity risk (Esterbauer et al., 2001). In the central nervous system, Ucp2 negatively regulates glucose-sensing in melanin-concentrating hormone expressing neurons in the lateral hypothalamus (Kong et al., 2010), a key region in the regulation of appetite. Ghrelin induced activation of neuropeptide Y/ agouti-related peptide neurons in the arcuate nucleus, another key pathway in the instigation of feeding behavior, are dependent on Ucp2 (Andrews et al., 2008), together indicating that genetic variants involved in energy regulation and appetite may extend to alter individual differences in responses to cues associated with food delivery. Ucp2, however, is likely involved in other processes as well, including regulation of anxiety-like behavior in mice (Hermes et al., 2016). The differential expression of Ucp2 in the PL and NAcc raises the possibility that heritable differences in the anticipatory response to palatable food are regulated by this gene system.

We found a cis-eQTL for *Wnt11* expression in the IL and BLA that colocalized with three behavioral QTLs, including lever presses during conditioned reinforcement. *Wnt11* has been previously implicated in acetylcholine and nicotinic receptor function. *Wnt* signaling is involved in nervous system development and may be involved in major psychiatric diseases such as schizophrenia and bipolar disorder (Okerlund & Cheyette, 2011). Interestingly, although *Wnt11* expression has been shown to enhance acetylcholine nicotinic receptor clustering in neuromuscular junctions (Messeant et al., 2017) it has heretofore not been identified in the CNS, so its relevance to the behaviors we examined is uncertain. Forebrain acetylcholine function has been identified as a major correlate differentiating sign- and goal-tracking (Paolone et al., 2013), where attentional top-down deficits in sign-trackers relative to goal-trackers appear to reflect attenuated cholinergic functioning in the basal forebrain (Kucinski et al., 2022; Paolone et al., 2013) involving choline transporter systems (Carmon et al., 2023). Further, work from our lab and others demonstrate that nicotinic receptor agonism facilitates sign-tracking (Overby et al., 2018; Palmatier et al., 2014; Palmatier et al., 2013; Versaggi et al., 2016), raising the intriguing possibility that central *Wnt11* may be involved in regulating the sign-tracking phenotype via CNS acetylcholine modulation. Interestingly, *Tenm4* loss of function in mice further impairs Wnt protein signaling (Nakamura et al., 2013) suggesting that these candidate genes we identified in our GWAS may interact to affect central nervous system function and behavior.

Finally, other cis-eQTLs identified genes in the central nervous system such as *Fchsd2* and *Fam168a* in high association to multiple traits, However, these known functions of these genes are limited by the lack of experimental behavioral studies using preclinical models examining function. *Fchsd2* and *Fam168a* have previously been identified as possible loci in human GWAS for nicotine dependence (Fchsd2; Gelernter et al., 2015) and smoking initiation (Fam168a; Liu et al., 2019). The lack of attention to these genes may make them attractive novel candidates for their role in complex behavior.

### Phenome-Wide Associations for Drug Response and Behavioral Regulation

We conducted a PheWAS to determine whether the genetic loci associated with the attribution of incentive salience in our study were also associated with other behavioral traits collected in other HS rat cohorts. Strikingly, a region identified on chromosome 1 (160 Mb) was strongly associated with acquisition of nicotine self-administration on the initial day of drug-taking (r^2^ = 0.84) and initial three sessions (r^2^=1). Regions on chromosome 1 were also more modestly associated with reinstatement to nicotine-seeking (Wang et al., 2018) (r^2^: 0.29 – 0.39), suggesting that genetic loci on chromosome 1 are pleiotropic in that they regulate the response to nicotine. We have previously shown that sign-trackers show heightened cue-induced reinstatement to nicotine-seeking in Sprague-Dawley rats (Versaggi et al., 2016), however until recently the genetic basis underlying these differences have not been characterized. Although there are likely many loci underlying the relationship between these two traits, these data suggest that this region of chromosome 1 contains key influential variants. It is notable that several of the genes identified here have been previously associated with features of smoking dependence in humans. We are currently conducting a separate GWAS in these HS rats for the genetic basis of socially-acquired adolescent nicotine self-administration, and future data will better identify the overlapping regions influencing these two traits.

In our cocaine sensitization paradigm however, PheWAS did not reveal associations that were strong as during nicotine self-administration. We found a modest (r^2^ = 0.19) association between a QTL identified for CRf operant responses on chromosome 2, and cocaine-induced head-waving on the final session of conditioning, but conditioned and unconditioned locomotion were largely unassociated (r^2^: 0.01-0.06). We have shown previously that sign- and goal-trackers do not show differences in locomotor response to a modest (10mg/kg i.p.) dose of cocaine, although sign-trackers show heightened unconditioned ultrasonic vocalizations (USVs) (Tripi et al., 2017). It is therefore perhaps not surprising that PheWAS yielded stronger associations with measures of nicotine response relative to cocaine, suggesting a different genetic relationship between sign-tracking and these two drug categories. However, cocaine self-administration involves processes other than locomotor sensitization. For example, sign-trackers are more sensitive to the presence of cocaine cues during drug-taking (Saunders & Robinson, 2010) and motivational properties of cocaine (Saunders & Robinson, 2011). We are currently testing a cohort of HS rats under several models of cocaine self-administration, including intermittent-access (Allain et al., 2018; Zimmer et al., 2012) and long-access self-administration (Lieselot et al., 2021), and ongoing work will determine whether the shared genetic basis of sign-tracking with cocaine responses depends on the model of cocaine conditioning.

Genetic loci identified for incentive salience attribution should theoretically be similar among separate cohorts of HS rats, even if the cohorts were tested at different locations, ages, and had different histories of behavioral testing. To test this, we used PheWAS to determine whether significant loci identified in our cohort at the University at Buffalo for PavCA would be associated with PavCA in another large cohort phenotyped at the University of Michigan (n = 1,583). The genetic correlations between both the Buffalo and Michigan cohorts were high (not shown). Notably, we found that a region reflected by a QTL on chromosome 18 (chr18: 85843691) that strongly influenced measures of goal-tracking and conditioned reinforcement (r^2^: 0.943-1) at both locations, although surprisingly there were no associations containing either chromosome 1, or measures of sign-tracking during PavCA. GWAS and PheWAS are therefore useful together for identifying loci across separate testing cohorts.

### Limitations and future directions

This study has several limitations. One caveat is that we have not presented sex-specific GWAS results due to power constraints. Tendency to sign-track is higher in females, a pattern that we also observed in this cohort of rats (King et al., 2021) and others (Hughson et al., 2019; King et al., 2016; Pitchers et al., 2015). To address this, we separately quantile normalized males and females before pooling them, which allowed us to avoid mean differences in tendency to sign- or goal-track across the different measures. We plan to continue to increase the size of our target PavCA population in the future by utilizing additional cohorts of HS and Sprague-Dawley rats for large-scale meta-analysis, which in the future will allow us to probe for sex-specific QTLs and gene-sex interactions.

A second limitation of this study is that the behavioral studies were conducted as part of a large multi-center project to assess the genetic and phenotypic relatedness of multiple addiction-associated traits in HS rats. Thus, the HS rats undergoing PavCA were not behaviorally naïve, but had instead undergone a battery of behavioral regulation tests before entering this study. As a result, the effect of certain genes on complex behavior may be influenced by age and testing history. We are working to address this in the future by including an additional cohort of HS rats tested at the University of Michigan who underwent PavCA prior to any behavioral testing. Genetic correlation (rG) values between the Buffalo and Michigan cohort are high (> 0.9) for terminal measures of PavCA, and by pooling these data we will be able to disentangle testing and age-related genetic effects.

## Conclusions

This study, using a large population of HS rats, identified multiple genes and loci associated with the attribution of incentive salience. Many of these genes are expressed in the central nervous system or have prior associations with psychiatric GWASs, and thus are attractive follow-up candidates for experimental behavioral genetics. Further, unlike traditional GWAS and omics studies for these traits in the domain of psychiatric disorders, these loci suggest roles in behavioral endophenotypes along a normal continuum of functioning in HS rats. This work supports the use of HS rats for the mapping of complex traits and provides several candidate genes for further additional studies on behavioral regulation.

## Supporting information

Supplementary Materials 1- GWAS Report Tables

Supplementary Materials 2- Manuscript Tables

Supplementary Materials 3 - GWAS Report

## Data Availability

The genotype data are available in the UC San Diego Digital Library Collection under the following citation: Palmer, Abraham A. (2023). “Heterogeneous Stock (HS) Rat Genotypes, Version 1. In Genotype Data from: NIDA Center for GWAS in Outbred Rats.” UC San Diego Library Digital Collections. https://doi.org/10.6075/J0028RR4.

## Dual Publication Statement

The primary findings, results, and conclusions presented in this paper address a different scientific question than those presented in King et al. 2021.

## Acknowledgements

Research reported in this publication was supported by grants from the National Institute on Alcohol Abuse and Alcoholism (AA024112 and AA007583), and the National Institute on Drug Abuse (P50 DA037844). The content is solely the responsibility of the authors and does not necessarily represent the official views of the NIH.

