## Supplementary Materials 3 - GWAS Report for "Genomic Loci Influencing Cue-Reactivity in Heterogeneous Stock Rats": Supplementary Materials 3 - GWAS Report.html

pavca\_crf\_report


### pavca\_crf\_report

##### ¶ Palmer Lab - UCSD ¶

###### 21 July, 2023

---

### 1 **Trait descriptions**

---

### 2 **Raw trait distribution**

#### 2.1 crf NY active inactive diff

#### 2.2 crf NY incentive value index

#### 2.3 crf NY lever presses

#### 2.4 pavca NY d5 avg lev lat

#### 2.5 pavca NY d5 avg mag lat

#### 2.6 pavca NY d5 index

#### 2.7 pavca NY d5 lever cs

#### 2.8 pavca NY d5 magazine cs

#### 2.9 pavca NY d5 magazine ncs

#### 2.10 pavca NY d5 prob lev

#### 2.11 pavca NY d5 prob mag

#### 2.12 pavca NY d5 response bias

---

### 3 **Phenotype processing details**

Quantile normalize separately for males and females then regress out
the effect of age/ cohort / box number if significant and explains more
than 2% of variance\*\*

```
Age not regressed out for any traits
```

- Following table lists the percent variance explained by other
  covariates

---

### 4 **SNP Heritability Estimates *h^2^***

---

### 5 **Summary of QTLs**

---

### 6 **Manhattan plots**

#### 6.1 crf NY active inactive diff

#### 6.2 crf NY incentive value index

#### 6.3 crf NY lever presses

  

| chr | number of qtls | topSNP | size of interval | topSNP -log10P |
| --- | --- | --- | --- | --- |
| chr1 | 1 | chr1:163339560 | 5.1 Mb | 9.529 |
| chr18 | 1 | chr18:85854393 | 0.65 Mb | 5.749 |
| chr2 | 1 | chr2:47800519 | 3 Mb | 5.612 |
| chr4 | 1 | chr4:135061594 | 2.4 Mb | 5.993 |

#### 6.4 pavca NY d5 avg lev lat

  

| chr | number of qtls | topSNP | size of interval | topSNP -log10P |
| --- | --- | --- | --- | --- |
| chr1 | 1 | chr1:159701681 | 1.8 Mb | 8.084 |
| chr18 | 1 | chr18:86488158 | 0.65 Mb | 5.852 |

#### 6.5 pavca NY d5 avg mag lat

#### 6.6 pavca NY d5 index

  

| chr | number of qtls | topSNP | size of interval | topSNP -log10P |
| --- | --- | --- | --- | --- |
| chr1 | 1 | chr1:160273674 | 1.9 Mb | 7.785 |
| chr18 | 1 | chr18:85843693 | 0.63 Mb | 7.123 |
| chr4 | 1 | chr4:133998243 | 2.9 Mb | 6.011 |

#### 6.7 pavca NY d5 lever cs

  

| chr | number of qtls | topSNP | size of interval | topSNP -log10P |
| --- | --- | --- | --- | --- |
| chr1 | 1 | chr1:160053871 | 1.8 Mb | 8.401 |
| chr18 | 1 | chr18:86488158 | 0.65 Mb | 6.254 |
| chr4 | 1 | chr4:69796826 | 0.53 Mb | 6.589 |

#### 6.8 pavca NY d5 magazine cs

  

| chr | number of qtls | topSNP | size of interval | topSNP -log10P |
| --- | --- | --- | --- | --- |
| chr1 | 1 | chr1:160448253 | 2.7 Mb | 6.787 |
| chr18 | 1 | chr18:85843693 | 0.63 Mb | 6.045 |

#### 6.9 pavca NY d5 magazine ncs

  

| chr | number of qtls | topSNP | size of interval | topSNP -log10P |
| --- | --- | --- | --- | --- |
| chr15 | 1 | chr15:74224438 | 2.7 Mb | 6.511 |

#### 6.10 pavca NY d5 prob lev

  

| chr | number of qtls | topSNP | size of interval | topSNP -log10P |
| --- | --- | --- | --- | --- |
| chr1 | 1 | chr1:166162182 | 4.6 Mb | 7.867 |
| chr18 | 1 | chr18:86488158 | 0.65 Mb | 6.391 |
| chr4 | 1 | chr4:69796826 | 0.53 Mb | 6.414 |

#### 6.11 pavca NY d5 prob mag

  

| chr | number of qtls | topSNP | size of interval | topSNP -log10P |
| --- | --- | --- | --- | --- |
| chr1 | 1 | chr1:167160047 | 1.3 Mb | 5.686 |

#### 6.12 pavca NY d5 response bias

  

| chr | number of qtls | topSNP | size of interval | topSNP -log10P |
| --- | --- | --- | --- | --- |
| chr1 | 1 | chr1:160204141 | 1.9 Mb | 9.225 |
| chr18 | 1 | chr18:85843693 | 0.63 Mb | 5.899 |
| chr4 | 1 | chr4:69793795 | 0.53 Mb | 6.555 |

---

### 7 **Regional Association plots**

[1] 26 [1] 26

#### 7.1 crf NY active inactive diff chr2:248891968

Size of interval : 77,254 bp

Number of genes in interval: 0

RGD link for genes in interval: NA  

##### 7.1.1 Putatively causal coding variants: crf NY active inactive diff chr2:248891968

HIGH or MODERATE impact variants absent

##### 7.1.2 eQTL info: crf NY active inactive diff chr2:248891968

No cis EQTLs detected

##### 7.1.3 PheWAS: P-values for other phenotypes at trait topSNP :

crf NY active inactive diff chr2:248891968

No pheWAS information

##### 7.1.4 PheWAS: Lowest P-values for other phenotypes in a 3Mb window

crf NY active inactive diff chr2:248891968

No pheWAS information

#### 7.2 crf NY incentive value index chr1:166077595

Size of interval : 4,633,990 bp

Number of genes in interval: 103

RGD link for genes in interval: Capn5
Omp
Arrb1
Mir326
Olr35
LOC102553078
Ppme1
LOC103691200
LOC102546541
Folr2
Rnf121
Rrm1
RGD1559902
Trnap-ugg1
Slco2b1
LOC102547456
Chrdl2
C2cd3
Mrpl48
Atg16l2
Art2b
Lamtor1
Lrrc32
Rps3
Tpbgl
Neu3
Pold3
Plekhb1
LOC108349453
Arap1
Clpb
LOC103691224
Klhl35
Ppid-ps5
Or2at1
Xrra1
Stard10
Ldha-ps3
Numa1
Il18bp
Art5
Nup98
Pgap2
Rps29-ps1
RGD1561870
Uvrag
Map6
Lipt2
Coa4
Mir3102
P2ry6
P2ry2
Fchsd2
Phox2a
Folr1
Chrna10
Gpx4-ps1
Rhog
Gdpd4
Acer3
Thap12
Mogat2
Olr36
Spcs2
Kcne3
Dnajb13
Relt
Mir139
LOC102547177
Anapc15
Lrtomt
Art1
Myo7a
Gucy2e
Wnt11
Dgat2
Serpinh1
Rnf169
LOC100912071
Pgm2l1
Ucp3
Ucp2
Rab6a
Fam168a
LOC103691225
Lrrc51
Xndc1
LOC108348605
Stim1
Or55b3
B3gnt6
Tsku
tuba1c-ps2
Emsy
Trnap-agg1
Gdpd5
Rpl23-ps3
P4ha3
Pde2a
Inppl1
Rpl12-ps3
Trpc2
Or55b4

#### 7.3 crf NY incentive value index chr2:248891968

Size of interval : 77,254 bp

Number of genes in interval: 0

RGD link for genes in interval: NA  

##### 7.3.1 Putatively causal coding variants: crf NY incentive value index chr2:248891968

HIGH or MODERATE impact variants absent

##### 7.3.2 eQTL info: crf NY incentive value index chr2:248891968

No cis EQTLs detected

##### 7.3.3 PheWAS: P-values for other phenotypes at trait topSNP :

crf NY incentive value index chr2:248891968

No pheWAS information

##### 7.3.4 PheWAS: Lowest P-values for other phenotypes in a 3Mb window

crf NY incentive value index chr2:248891968

No pheWAS information

#### 7.4 crf NY incentive value index chr4:135061594

Size of interval : 2,426,565 bp

Number of genes in interval: 8

RGD link for genes in interval: Smtn-ps1
LOC108350739
Pdzrn3
LOC102555697
Cntn3
LOC108350740
Tuba1b-ps2
Rpl13-ps14

#### 7.5 crf NY lever presses chr18:85854393

Size of interval : 649,463 bp

Number of genes in interval: 9

RGD link for genes in interval: Pclaf-ps2
Rttn
Socs6
Chn3
LOC689166
Cd226
Dok6
LOC100362807
Snapc5-ps1  

##### 7.5.1 Putatively causal coding variants: crf NY lever presses chr18:85854393

HIGH or MODERATE impact variants absent

##### 7.5.2 eQTL info: crf NY lever presses chr18:85854393

No cis EQTLs detected

##### 7.5.3 PheWAS: P-values for other phenotypes at trait topSNP :

crf NY lever presses chr18:85854393

No pheWAS information

##### 7.5.4 PheWAS: Lowest P-values for other phenotypes in a 3Mb window

crf NY lever presses chr18:85854393

No pheWAS information

#### 7.6 crf NY lever presses chr1:163339560

Size of interval : 5,127,829 bp

Number of genes in interval: 113

RGD link for genes in interval: Hmgb1-ps23
Capn5
Omp
Arrb1
Mir326
Olr35
LOC102553078
Ppme1
LOC103691200
LOC102546541
Folr2
Rnf121
Rrm1
Rsf1
RGD1559902
Trnap-ugg1
Slco2b1
LOC102547456
Chrdl2
C2cd3
Mrpl48
Atg16l2
Art2b
Lamtor1
Clns1a
Aqp11
Pak1
Lrrc32
Rps3
Tpbgl
Neu3
Pold3
Plekhb1
LOC108349453
Arap1
Clpb
LOC103691224
Ints4
Klhl35
Ppid-ps5
Or2at1
Xrra1
Stard10
Ldha-ps3
Numa1
Il18bp
Art5
Nup98
Pgap2
Rps29-ps1
Pkm-ps7
RGD1561870
Uvrag
Map6
Lipt2
Coa4
Mir3102
P2ry6
P2ry2
Fchsd2
Phox2a
Folr1
Chrna10
Gpx4-ps1
Rhog
Aamdc
Gdpd4
Acer3
Thap12
Mogat2
Olr36
Spcs2
Kcne3
Dnajb13
Relt
Mir139
LOC102547177
Anapc15
Lrtomt
Art1
Rpl35-ps4
Myo7a
Gucy2e
Wnt11
Dgat2
Serpinh1
Rnf169
LOC100912071
Pgm2l1
Ucp3
Ucp2
Rab6a
Fam168a
LOC103691225
Lrrc51
Xndc1
LOC108348605
Stim1
Or55b3
LOC102550562
B3gnt6
Tsku
tuba1c-ps2
Emsy
Trnap-agg1
Gdpd5
Rpl23-ps3
P4ha3
Pde2a
Inppl1
Rpl12-ps3
Trpc2
Or55b4

#### 7.7 crf NY lever presses chr2:47800519

Size of interval : 2,960,705 bp

Number of genes in interval: 14

RGD link for genes in interval: Arl15
Or8b54c
LOC103691459
Itga2
Pelo
LOC103691460
LOC108350167
Mfap1a-ps1
Ndufs4
Fst
Mocs2
Rpl29-ps16
Itga1
Isl1  

##### 7.7.1 Putatively causal coding variants: crf NY lever presses chr2:47800519

HIGH or MODERATE impact variants absent

##### 7.7.2 eQTL info: crf NY lever presses chr2:47800519

Trait topSNP : chr2:47800519

Trait topSNP -log10P : 5.612

###### 7.7.2.1 Itga2

Ensembl gene name : ENSRNOG00000058111

***integrin alpha 2***

*RGD link:* Itga2

*Human GWAS Catalog link:* Itga2

*Pubmed link:* Itga2

*Alliance of Genome Resources link:* Itga2

###### 7.7.2.2 Mocs2

Ensembl gene name : ENSRNOG00000056325

***molybdenum cofactor synthesis 2***

*RGD link:* Mocs2

*Human GWAS Catalog link:* *No Human GWAS Catalog
entry*

*Pubmed link:* Mocs2

*Alliance of Genome Resources link:* Mocs2

###### 7.7.2.3 Pelo

Ensembl gene name : ENSRNOG00000061128

***pelota homolog (Drosophila)***

*RGD link:* Pelo

*Human GWAS Catalog link:* Pelo

*Pubmed link:* Pelo

*Alliance of Genome Resources link:* Pelo

##### 7.7.3 PheWAS: P-values for other phenotypes at trait topSNP :

crf NY lever presses chr2:47800519

No pheWAS information

##### 7.7.4 PheWAS: Lowest P-values for other phenotypes in a 3Mb window

crf NY lever presses chr2:47800519

No pheWAS information

#### 7.8 crf NY lever presses chr4:135061594

Size of interval : 2,426,565 bp

Number of genes in interval: 8

RGD link for genes in interval: Smtn-ps1
LOC108350739
Pdzrn3
LOC102555697
Cntn3
LOC108350740
Tuba1b-ps2
Rpl13-ps14  

##### 7.8.1 Putatively causal coding variants: crf NY lever presses chr4:135061594

HIGH or MODERATE impact variants absent

##### 7.8.2 eQTL info: crf NY lever presses chr4:135061594

Trait topSNP : chr4:135061594

Trait topSNP -log10P : 5.993

###### 7.8.2.1 Cntn3

Ensembl gene name : ENSRNOG00000006144

***contactin 3***

*RGD link:* Cntn3

*Human GWAS Catalog link:* Cntn3

*Pubmed link:* Cntn3

*Alliance of Genome Resources link:* Cntn3

###### 7.8.2.2 Pdzrn3

Ensembl gene name : ENSRNOG00000057556

***PDZ domain containing RING finger 3***

*RGD link:* Pdzrn3

*Human GWAS Catalog link:* Pdzrn3

*Pubmed link:* Pdzrn3

*Alliance of Genome Resources link:* Pdzrn3

##### 7.8.3 PheWAS: P-values for other phenotypes at trait topSNP :

crf NY lever presses chr4:135061594

No pheWAS information

##### 7.8.4 PheWAS: Lowest P-values for other phenotypes in a 3Mb window

crf NY lever presses chr4:135061594

No pheWAS information

#### 7.9 pavca NY d5 avg lev lat chr18:86488158

Size of interval : 649,463 bp

Number of genes in interval: 9

RGD link for genes in interval: Pclaf-ps2
Rttn
Socs6
Chn3
LOC689166
Cd226
Dok6
LOC100362807
Snapc5-ps1  

##### 7.9.1 Putatively causal coding variants: pavca NY d5 avg lev lat chr18:86488158

HIGH or MODERATE impact variants absent

##### 7.9.2 eQTL info: pavca NY d5 avg lev lat chr18:86488158

No cis EQTLs detected

##### 7.9.3 PheWAS: P-values for other phenotypes at trait topSNP :

pavca NY d5 avg lev lat chr18:86488158

No pheWAS information

##### 7.9.4 PheWAS: Lowest P-values for other phenotypes in a 3Mb window

pavca NY d5 avg lev lat chr18:86488158

No pheWAS information

#### 7.10 pavca NY d5 avg lev lat chr1:159701681

Size of interval : 1,786,021 bp

Number of genes in interval: 4

RGD link for genes in interval: Tenm4
Pdzd11-ps1
Mir708
Rpl23a-ps7

#### 7.11 pavca NY d5 index chr18:85843693

Size of interval : 630,283 bp

Number of genes in interval: 9

RGD link for genes in interval: Pclaf-ps2
Rttn
Socs6
Chn3
LOC689166
Cd226
Dok6
LOC100362807
Snapc5-ps1

#### 7.12 pavca NY d5 index chr1:160273674

Size of interval : 1,876,718 bp

Number of genes in interval: 4

RGD link for genes in interval: Tenm4
Pdzd11-ps1
Mir708
Rpl23a-ps7

#### 7.13 pavca NY d5 index chr4:133998243

Size of interval : 2,910,048 bp

Number of genes in interval: 12

RGD link for genes in interval: Gxylt2
Smtn-ps1
Ppp4r2
LOC108350739
Pdzrn3
LOC689383
Shq1
LOC102555697
Cntn3
LOC108350740
Tuba1b-ps2
Rpl13-ps14

#### 7.14 pavca NY d5 lever cs chr18:86488158

Size of interval : 649,463 bp

Number of genes in interval: 9

RGD link for genes in interval: Pclaf-ps2
Rttn
Socs6
Chn3
LOC689166
Cd226
Dok6
LOC100362807
Snapc5-ps1  

##### 7.14.1 Putatively causal coding variants: pavca NY d5 lever cs chr18:86488158

HIGH or MODERATE impact variants absent

##### 7.14.2 eQTL info: pavca NY d5 lever cs chr18:86488158

No cis EQTLs detected

##### 7.14.3 PheWAS: P-values for other phenotypes at trait topSNP :

pavca NY d5 lever cs chr18:86488158

No pheWAS information

##### 7.14.4 PheWAS: Lowest P-values for other phenotypes in a 3Mb window

pavca NY d5 lever cs chr18:86488158

No pheWAS information

#### 7.15 pavca NY d5 lever cs chr1:160053871

Size of interval : 1,786,021 bp

Number of genes in interval: 4

RGD link for genes in interval: Tenm4
Pdzd11-ps1
Mir708
Rpl23a-ps7

#### 7.16 pavca NY d5 lever cs chr4:69796826

Size of interval : 528,398 bp

Number of genes in interval: 2

RGD link for genes in interval: Rpl30-ps10
Rpl4-ps2

#### 7.17 pavca NY d5 magazine cs chr18:85843693

Size of interval : 630,283 bp

Number of genes in interval: 9

RGD link for genes in interval: Pclaf-ps2
Rttn
Socs6
Chn3
LOC689166
Cd226
Dok6
LOC100362807
Snapc5-ps1  

##### 7.17.1 Putatively causal coding variants: pavca NY d5 magazine cs chr18:85843693

HIGH or MODERATE impact variants absent

##### 7.17.2 eQTL info: pavca NY d5 magazine cs chr18:85843693

No cis EQTLs detected

##### 7.17.3 PheWAS: P-values for other phenotypes at trait topSNP :

pavca NY d5 magazine cs chr18:85843693

No pheWAS information

##### 7.17.4 PheWAS: Lowest P-values for other phenotypes in a 3Mb window

pavca NY d5 magazine cs chr18:85843693

No pheWAS information

#### 7.18 pavca NY d5 magazine cs chr1:160448253

Size of interval : 2,691,650 bp

Number of genes in interval: 23

RGD link for genes in interval: Hmgb1-ps23
Tenm4
Gab2
Prdx1-ps2
Rsf1
Clns1a
Aqp11
Pak1
Mir708
LOC102550214
Ints4
Usp35
Alg8
Kctd21
Ndufc2
Thrsp
Aamdc
Rpl23a-ps7
Kctd14
Rpl35-ps4
Nars2
LOC108349194
LOC102550562

#### 7.19 pavca NY d5 magazine ncs chr15:74224438

Size of interval : 2,702,701 bp

Number of genes in interval: 8

RGD link for genes in interval: Nono-ps9
Mtfr2-ps1
Envl-ps1
LOC102547864
LOC102546653
Cdk4-ps1
LOC498555
Phgdh-ps4  

##### 7.19.1 Putatively causal coding variants: pavca NY d5 magazine ncs chr15:74224438

HIGH or MODERATE impact variants absent

##### 7.19.2 eQTL info: pavca NY d5 magazine ncs chr15:74224438

No cis EQTLs detected

##### 7.19.3 PheWAS: P-values for other phenotypes at trait topSNP :

pavca NY d5 magazine ncs chr15:74224438

No pheWAS information

##### 7.19.4 PheWAS: Lowest P-values for other phenotypes in a 3Mb window

pavca NY d5 magazine ncs chr15:74224438

No pheWAS information

#### 7.20 pavca NY d5 prob lev chr4:69796826

Size of interval : 528,398 bp

Number of genes in interval: 2

RGD link for genes in interval: Rpl30-ps10
Rpl4-ps2

#### 7.21 pavca NY d5 prob lev chr1:166162182

Size of interval : 4,633,990 bp

Number of genes in interval: 103

RGD link for genes in interval: Capn5
Omp
Arrb1
Mir326
Olr35
LOC102553078
Ppme1
LOC103691200
LOC102546541
Folr2
Rnf121
Rrm1
RGD1559902
Trnap-ugg1
Slco2b1
LOC102547456
Chrdl2
C2cd3
Mrpl48
Atg16l2
Art2b
Lamtor1
Lrrc32
Rps3
Tpbgl
Neu3
Pold3
Plekhb1
LOC108349453
Arap1
Clpb
LOC103691224
Klhl35
Ppid-ps5
Or2at1
Xrra1
Stard10
Ldha-ps3
Numa1
Il18bp
Art5
Nup98
Pgap2
Rps29-ps1
RGD1561870
Uvrag
Map6
Lipt2
Coa4
Mir3102
P2ry6
P2ry2
Fchsd2
Phox2a
Folr1
Chrna10
Gpx4-ps1
Rhog
Gdpd4
Acer3
Thap12
Mogat2
Olr36
Spcs2
Kcne3
Dnajb13
Relt
Mir139
LOC102547177
Anapc15
Lrtomt
Art1
Myo7a
Gucy2e
Wnt11
Dgat2
Serpinh1
Rnf169
LOC100912071
Pgm2l1
Ucp3
Ucp2
Rab6a
Fam168a
LOC103691225
Lrrc51
Xndc1
LOC108348605
Stim1
Or55b3
B3gnt6
Tsku
tuba1c-ps2
Emsy
Trnap-agg1
Gdpd5
Rpl23-ps3
P4ha3
Pde2a
Inppl1
Rpl12-ps3
Trpc2
Or55b4  

##### 7.21.1 Putatively causal coding variants: pavca NY d5 prob lev chr1:166162182

##### 7.21.2 eQTL info: pavca NY d5 prob lev chr1:166162182

Trait topSNP : chr1:166162182

Trait topSNP -log10P : 7.867

###### 7.21.2.1 AABR07004876.1

Ensembl gene name : ENSRNOG00000018083

*RGD link:* *No RGD entry*

###### 7.21.2.2 Acer3

Ensembl gene name : ENSRNOG00000036866

***alkaline ceramidase 3***

*RGD link:* Acer3

*Human GWAS Catalog link:* Acer3

*Pubmed link:* Acer3

*Alliance of Genome Resources link:* Acer3

###### 7.21.2.3 Arap1

Ensembl gene name : ENSRNOG00000019555

***ArfGAP with RhoGAP domain, ankyrin repeat and PH domain
1***

*RGD link:* Arap1

*Human GWAS Catalog link:* Arap1

*Pubmed link:* Arap1

*Alliance of Genome Resources link:* Arap1

###### 7.21.2.4 Atg16l2

Ensembl gene name : ENSRNOG00000019413

***autophagy related 16-like 2***

*RGD link:* Atg16l2

*Human GWAS Catalog link:* Atg16l2

*Pubmed link:* Atg16l2

*Alliance of Genome Resources link:* Atg16l2

###### 7.21.2.5 Capn5

Ensembl gene name : ENSRNOG00000014251

***calpain 5***

*RGD link:* Capn5

*Human GWAS Catalog link:* Capn5

*Pubmed link:* Capn5

*Alliance of Genome Resources link:* Capn5

###### 7.21.2.6 Dgat2

Ensembl gene name : ENSRNOG00000016573

***diacylglycerol O-acyltransferase 2***

*RGD link:* Dgat2

*Human GWAS Catalog link:* Dgat2

*Pubmed link:* Dgat2

*Alliance of Genome Resources link:* Dgat2

###### 7.21.2.7 Fam168a

Ensembl gene name : ENSRNOG00000018873

***family with sequence similarity 168, member A***

*RGD link:* Fam168a

*Human GWAS Catalog link:* Fam168a

*Pubmed link:* Fam168a

*Alliance of Genome Resources link:* Fam168a

###### 7.21.2.8 Fchsd2

Ensembl gene name : ENSRNOG00000019319

***FCH and double SH3 domains 2***

*RGD link:* Fchsd2

*Human GWAS Catalog link:* Fchsd2

*Pubmed link:* Fchsd2

*Alliance of Genome Resources link:* Fchsd2

###### 7.21.2.9 Folr1

Ensembl gene name : ENSRNOG00000019902

***folate receptor 1***

*RGD link:* Folr1

*Human GWAS Catalog link:* *No Human GWAS Catalog
entry*

*Pubmed link:* Folr1

*Alliance of Genome Resources link:* Folr1

###### 7.21.2.10 Gucy2e

Ensembl gene name : ENSRNOG00000015058

***guanylate cyclase 2E***

*RGD link:* Gucy2e

*Human GWAS Catalog link:* *No Human GWAS Catalog
entry*

*Pubmed link:* *NA*

*Alliance of Genome Resources link:* *NA*

###### 7.21.2.11 Lipt2

Ensembl gene name : ENSRNOG00000016906

***lipoyl(octanoyl) transferase 2 (putative)***

*RGD link:* Lipt2

*Human GWAS Catalog link:* *No Human GWAS Catalog
entry*

*Pubmed link:* Lipt2

*Alliance of Genome Resources link:* Lipt2

###### 7.21.2.12 Map6

Ensembl gene name : ENSRNOG00000027204

***microtubule-associated protein 6***

*RGD link:* Map6

*Human GWAS Catalog link:* *No Human GWAS Catalog
entry*

*Pubmed link:* Map6

*Alliance of Genome Resources link:* Map6

###### 7.21.2.13 Mogat2

Ensembl gene name : ENSRNOG00000027228

***monoacylglycerol O-acyltransferase 2***

*RGD link:* Mogat2

*Human GWAS Catalog link:* Mogat2

*Pubmed link:* Mogat2

*Alliance of Genome Resources link:* Mogat2

###### 7.21.2.14 Numa1

Ensembl gene name : ENSRNOG00000000417

***nuclear mitotic apparatus protein 1***

*RGD link:* Numa1

*Human GWAS Catalog link:* Numa1

*Pubmed link:* Numa1

*Alliance of Genome Resources link:* Numa1

###### 7.21.2.15 Nup98

Ensembl gene name : ENSRNOG00000020347

***nucleoporin 98***

*RGD link:* Nup98

*Human GWAS Catalog link:* Nup98

*Pubmed link:* Nup98

*Alliance of Genome Resources link:* Nup98

###### 7.21.2.16 Pak1

Ensembl gene name : ENSRNOG00000029784

***p21 (RAC1) activated kinase 1***

*RGD link:* Pak1

*Human GWAS Catalog link:* Pak1

*Pubmed link:* Pak1

*Alliance of Genome Resources link:* Pak1

###### 7.21.2.17 Pgap2

Ensembl gene name : ENSRNOG00000020371

***post-GPI attachment to proteins 2***

*RGD link:* Pgap2

*Human GWAS Catalog link:* *No Human GWAS Catalog
entry*

*Pubmed link:* Pgap2

*Alliance of Genome Resources link:* Pgap2

###### 7.21.2.18 Pgm2l1

Ensembl gene name : ENSRNOG00000017079

***phosphoglucomutase 2-like 1***

*RGD link:* Pgm2l1

*Human GWAS Catalog link:* Pgm2l1

*Pubmed link:* Pgm2l1

*Alliance of Genome Resources link:* Pgm2l1

###### 7.21.2.19 Ppme1

Ensembl gene name : ENSRNOG00000017227

***protein phosphatase methylesterase 1***

*RGD link:* Ppme1

*Human GWAS Catalog link:* *No Human GWAS Catalog
entry*

*Pubmed link:* Ppme1

*Alliance of Genome Resources link:* Ppme1

###### 7.21.2.20 Rnf121

Ensembl gene name : ENSRNOG00000020175

***ring finger protein 121***

*RGD link:* Rnf121

*Human GWAS Catalog link:* *No Human GWAS Catalog
entry*

*Pubmed link:* Rnf121

*Alliance of Genome Resources link:* Rnf121

###### 7.21.2.21 Rps3

Ensembl gene name : ENSRNOG00000017418

***ribosomal protein S3***

*RGD link:* Rps3

*Human GWAS Catalog link:* Rps3

*Pubmed link:* Rps3

*Alliance of Genome Resources link:* Rps3

###### 7.21.2.22 Serpinh1

Ensembl gene name : ENSRNOG00000016831

***serpin family H member 1***

*RGD link:* Serpinh1

*Human GWAS Catalog link:* Serpinh1

*Pubmed link:* Serpinh1

*Alliance of Genome Resources link:* Serpinh1

###### 7.21.2.23 Slco2b1

Ensembl gene name : ENSRNOG00000017976

***solute carrier organic anion transporter family, member
2b1***

*RGD link:* Slco2b1

*Human GWAS Catalog link:* *No Human GWAS Catalog
entry*

*Pubmed link:* Slco2b1

*Alliance of Genome Resources link:* Slco2b1

###### 7.21.2.24 Ucp2

Ensembl gene name : ENSRNOG00000017854

***uncoupling protein 2***

*RGD link:* Ucp2

*Human GWAS Catalog link:* *No Human GWAS Catalog
entry*

*Pubmed link:* Ucp2

*Alliance of Genome Resources link:* Ucp2

###### 7.21.2.25 Uvrag

Ensembl gene name : ENSRNOG00000016206

***UV radiation resistance associated***

*RGD link:* Uvrag

*Human GWAS Catalog link:* Uvrag

*Pubmed link:* Uvrag

*Alliance of Genome Resources link:* Uvrag

###### 7.21.2.26 Wnt11

Ensembl gene name : ENSRNOG00000015982

***wingless-type MMTV integration site family, member
11***

*RGD link:* Wnt11

*Human GWAS Catalog link:* Wnt11

*Pubmed link:* Wnt11

*Alliance of Genome Resources link:* Wnt11

##### 7.21.3 PheWAS: P-values for other phenotypes at trait topSNP :

pavca NY d5 prob lev chr1:166162182

No pheWAS information

##### 7.21.4 PheWAS: Lowest P-values for other phenotypes in a 3Mb window

pavca NY d5 prob lev chr1:166162182

No pheWAS information

#### 7.22 pavca NY d5 prob lev chr18:86488158

Size of interval : 649,463 bp

Number of genes in interval: 9

RGD link for genes in interval: Pclaf-ps2
Rttn
Socs6
Chn3
LOC689166
Cd226
Dok6
LOC100362807
Snapc5-ps1  

##### 7.22.1 Putatively causal coding variants: pavca NY d5 prob lev chr18:86488158

HIGH or MODERATE impact variants absent

##### 7.22.2 eQTL info: pavca NY d5 prob lev chr18:86488158

No cis EQTLs detected

##### 7.22.3 PheWAS: P-values for other phenotypes at trait topSNP :

pavca NY d5 prob lev chr18:86488158

No pheWAS information

##### 7.22.4 PheWAS: Lowest P-values for other phenotypes in a 3Mb window

pavca NY d5 prob lev chr18:86488158

No pheWAS information

#### 7.23 pavca NY d5 prob mag chr1:167160047

Size of interval : 1,263,776 bp

Number of genes in interval: 34

RGD link for genes in interval: LOC102546541
Folr2
Rnf121
Atg16l2
Art2b
Lamtor1
Arap1
Clpb
Stard10
Ldha-ps3
Numa1
Il18bp
Art5
Nup98
Pgap2
Fchsd2
Phox2a
Folr1
Chrna10
Gpx4-ps1
Rhog
Mir139
LOC102547177
Anapc15
Lrtomt
Art1
LOC103691225
Lrrc51
Xndc1
LOC108348605
Pde2a
Inppl1
Rpl12-ps3
Trpc2

#### 7.24 pavca NY d5 response bias chr4:69793795

Size of interval : 528,398 bp

Number of genes in interval: 2

RGD link for genes in interval: Rpl30-ps10
Rpl4-ps2  

##### 7.24.1 Putatively causal coding variants: pavca NY d5 response bias chr4:69793795

HIGH or MODERATE impact variants absent

##### 7.24.2 eQTL info: pavca NY d5 response bias chr4:69793795

Trait topSNP : chr4:69793795

Trait topSNP -log10P : 6.555

###### 7.24.2.1 Chl1

Ensembl gene name : ENSRNOG00000045771

***cell adhesion molecule L1-like***

*RGD link:* Chl1

*Human GWAS Catalog link:* Chl1

*Pubmed link:* Chl1

*Alliance of Genome Resources link:* Chl1

##### 7.24.3 PheWAS: P-values for other phenotypes at trait topSNP :

pavca NY d5 response bias chr4:69793795

No pheWAS information

##### 7.24.4 PheWAS: Lowest P-values for other phenotypes in a 3Mb window

pavca NY d5 response bias chr4:69793795

No pheWAS information

#### 7.25 pavca NY d5 response bias chr1:160204141

Size of interval : 1,930,409 bp

Number of genes in interval: 4

RGD link for genes in interval: Tenm4
Pdzd11-ps1
Mir708
Rpl23a-ps7  

##### 7.25.1 Putatively causal coding variants: pavca NY d5 response bias chr1:160204141

HIGH or MODERATE impact variants absent

##### 7.25.2 eQTL info: pavca NY d5 response bias chr1:160204141

No cis EQTLs detected

##### 7.25.3 PheWAS: P-values for other phenotypes at trait topSNP :

pavca NY d5 response bias chr1:160204141

No pheWAS information

##### 7.25.4 PheWAS: Lowest P-values for other phenotypes in a 3Mb window

pavca NY d5 response bias chr1:160204141

No pheWAS information

#### 7.26 pavca NY d5 response bias chr18:85843693

Size of interval : 630,283 bp

Number of genes in interval: 9

RGD link for genes in interval: Pclaf-ps2
Rttn
Socs6
Chn3
LOC689166
Cd226
Dok6
LOC100362807
Snapc5-ps1
